# Contractility governs cardiomyocyte cell cycle activity

**DOI:** 10.1101/2025.08.19.671096

**Authors:** Nancy Shehata, Rajiven Srikantharajah, Eleonore Baier, Christoph Manthey, Tim Stüdemann, Marie Nehring, Anita Covic, Christian Müller, Bente Siebels, Paula Nissen, Anika Witten, Magdalena Zafra Castellano, Yago Rodriguez, Lucia Zamacona Gomez, Imke Janssen, Jorge Rodriguez, Andrea Garza Enriquez, Miguel Angel Garitagoitia Romero, Romina Di Mattia, Andreas Dendorfer, Alessandra Moretti, Christine Maria Poch, Thomas Eschenhagen, Florian Weinberger

## Abstract

The heart’s rhythmic contractions keep blood flowing to deliver the oxygen and nutrients that sustain life, but this very strength may also explain why the heart has so little capacity to regenerate. Whereas embryonic and neonatal hearts can repair themselves, this ability is rapidly lost after birth as heart muscle cells (cardiomyocytes) exit the cell cycle. We tested whether the force-generating machinery of the heart acts as a barrier to regeneration. Across human cardiomyocytes, engineered heart tissue, and myocardial slices, we found that active force generation suppresses proliferation, whereas its inhibition, through chemogenetic or pharmacological modulation, enabled cell cycle re-entry. These findings identify contractility as a central regulator of regenerative potential, highlighting new directions for strategies to restore repair in the diseased heart.

## Introduction

The primary function of the heart is to contract rhythmically to pump blood throughout the body, delivering oxygen and nutrients. This task is carried out by cardiomyocytes, highly specialized muscle cells that beat at a rate of 60-80 times per minute. While continuous contractile activity is essential for life, it may paradoxically limit the heart’s ability to regenerate. During embryonic development and early after birth the heart exhibits a remarkable regenerative potential (^1,2^). Loss of up to 60% of cardiomyocytes can be compensated during embryonic development (^1^). Likewise, profound regenerative responses have been observed in the neonatal heart, with functional recovery following apical resection or myocardial infarction in both mice and pigs (^3,4^). Limited clinical and pathological observations suggest that human neonates may also exhibit a brief postnatal window of regenerative capacity (^5–7^). This regenerative ability is lost within the first few days after birth as cardiomyocytes exit the cell cycle and heart growth occurs predominantly from cardiomyocyte hypertrophy (^8^). Cardiomyocyte loss can occur acutely, e.g. myocardial infarction) or gradually over time (e.g. long-term hypertension, myocarditis, or structural heart disease (^9–11^). Ultimately, heart failure represents a common sequela, the progression of which can be mitigated, and life expectancy improved through guideline-directed combination therapy, but the underlying damage cannot be reversed (^12^).

Cardiomyocyte contractile function is driven by sarcomeres, highly ordered protein complexes that constitute more than 50% of cardiomyocyte volume (^13–15^). Active force generation depends on the cyclic interactions between myosin and actin, but more than 100 proteins contribute to sarcomere function (^16^). While essential for maintaining cardiac contractility, sarcomeres form a rigid physical barrier to cell division, potentially limiting cardiomyocyte proliferation. The hypothesis that contractility restricts cardiomyocyte proliferation is supported by studies in patients receiving left-ventricular assist devices (LVADs), where mechanical unloading correlated with increased cardiomyocyte cell cycle activity, indicating the presence of a latent regenerative capacity (^17–19^). Additional evidence linking contractility, sarcomere structure, and cell cycle regulation comes from genetic and pharmacological interventions designed to induce cardiomyocyte proliferation. These approaches consistently led to sarcomere disassembly, accompanied by a transient decline in contractility or overall cardiac function in mice or stem cell-derived cardiac tissues (^20–26^). Together, these findings support the hypothesis that continuous mechanical demand suppresses cardiomyocyte cell cycle re-entry, a concept termed "All work and no repair” (^27^). To directly investigate this relationship, we employed both chemogenetic and pharmacological strategies to modulate cardiac contractility with high spatial and temporal precision. Inhibition of active force generation, with either chemogenetics or the cardiac myosin modulator mavacamten, increased cell cycle activity in human induced pluripotent stem cell (hiPSC)-derived cardiomyocytes, engineered heart tissue, and in human cardiac slices. Restoration of contractility induced cardiomyocytes to exit the cell cycle, providing strong evidence for a causal link between contractility and cardiomyocyte cell cycle arrest.

## Results

### Inhibition of contractility triggered sarcomere disassembly

To dissect the role of contractility in regulating cell cycle activity, we combined chemogenetic tools that enable rapid, reversible, and targeted modulation of cardiomyocyte contractility with pharmacological strategies **(Fig. 1A)**. Chemogenetic tools offer precise control with minimal off-target effects. Specifically, we employed PSAM-GlyR and PSAM^4^-GlyR to modulate contractile activity in a temporally controlled manner (^28–31^. HiPSC-derived cardiomyocytes expressing PSAM-GlyR were used to cast engineered heart tissue (EHTs). PSAM-GlyR EHTs exhibited immediate and complete cessation of contractile activity upon application of the agonist PSEM^89S^, due to a depolarization block (^29^). Importantly, this reversible contractility off-switch did not compromise cell viability, as evidenced by stable LDH release, unchanged TUNEL staining, a decrease in troponin I concentration in the medium and full recovery of contractile force post-washout **(Fig. S1 and Movie S1)**. To assess structural consequences of contractility inhibition, we analyzed sarcomere organization in EHTs cultured for 28 or 70 days respectively. Immunostaining for the Z-disk protein α-actinin revealed disorganized sarcomere structure in EHTs, in which contractility was inhibited for 7 days. Additional staining with phalloidin (I-band) and anti-titin antibodies (C-terminal M8-M9-M10; M-band) confirmed these findings. Quantitative analysis was performed with SarcAsM on alpha-actinin stained sections, which turned out to be the most sensitive parameter to detect even subtle changes in sarcomere structure (9-12 EHTs from three different cardiomyocyte batches; **Fig. 1B-D**). Taking advantage of the reversible nature of the chemogenetic system, we analyzed sarcomere architecture following re-initiation of contractility. Consistent with functional recovery, histological analysis confirmed restoration of sarcomere organization **(Fig. 1B)**. Beyond cytoskeletal changes, contractility inhibition also significantly affected nuclear morphology. Specifically, cardiomyocyte nuclei, which typically elongate over time in 3D EHT culture, adopted a more rounded morphology during contractility suppression, as shown by a significant increase in nuclear circularity (9-12 EHTs from three different cardiomyocyte batches; **Fig. 1B, E and F**).

**Figure 1:**
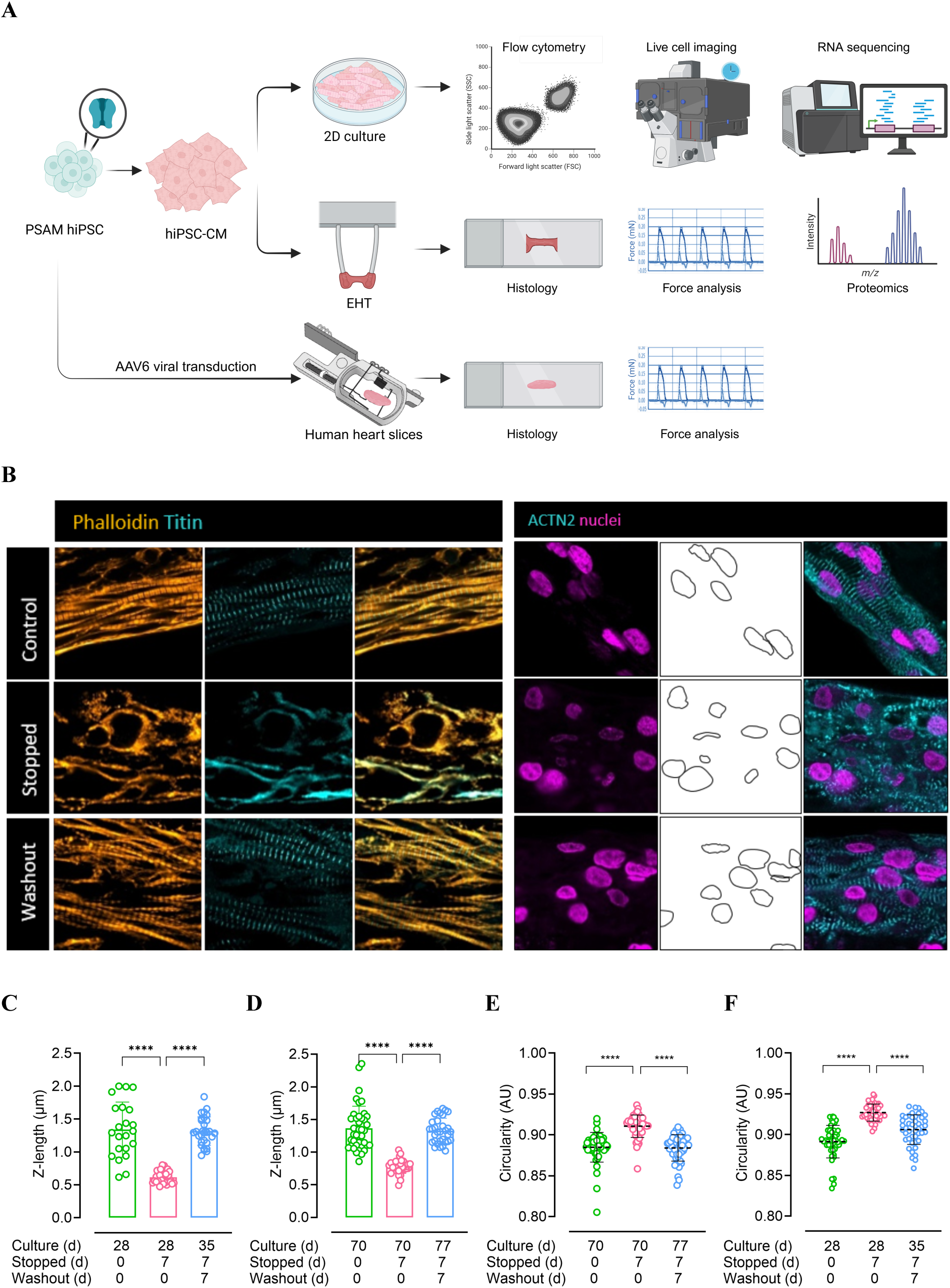
Contractility inhibition induced reversible cardiomyocyte remodelling. **(A)** Schematic depiction of the experimental set-up. (**B)** Contractility inhibition (7 days) triggered sarcomere disassembly, evidenced by immunostaining for actin (I-band), titin (M-band), and α-actinin (Z-disc), accompanied by increased nuclear circularity in cardiomyocytes. (**C-D)** Quantification of sarcomere organization using SarcAsM (n = 9). (**E-F)** Measurement of nuclear circularity using Stardist in beating, stopped and recovered EHTs cultured for 21 or 70 days (n = 9). Statistical analysis was performed using ANOVA. Significance levels are indicated as p < 0.00001 (****).

### Contractility inhibition promoted cardiomyocyte proliferation

We next analyzed the consequence of contractility inhibition on cell cycle activity in cardiomyocytes cultured in a 2D format. Contractility was inhibited by applying the nicotinic receptor agonist varenicline (100 nM) to PSAM^4^-GlyR cardiomyocytes for 7 days. DNA synthesis was assessed by EdU incorporation (**Fig. 2A**). Inhibition of contractility resulted in a significant increase in EdU-positive cardiomyocytes from 5.0% ± 0.5 to 14.3% ± 0.6 (n=3 experiments / N=3 biological replicates per experiment; **Fig. 2B-C**). Upon re-initiation of contractility EdU incorporation declined to 3.0% ± 0.4. To distinguish true proliferation (i.e. cytokinesis) from polyploidization, we assessed nuclear ploidy. This analysis revealed that inhibiting contractility stimulated overall cell cycle activity, as indicated by an increase in 4n cardiomyocytes, either binucleate diploid cells or mononuclear tetraploid cells. Importantly, there was also a significant increase in diploid, mononuclear EdU-positive cardiomyocytes (3.8 ± 1.5% vs. 1.4 ± 1.0% in beating controls), providing evidence for an increase in true cardiomyocyte proliferation (**Fig. 2D-E**). Further supporting this finding, live-cell imaging captured complete cardiomyocyte division events, characterized by chromosome segregation and cytokinesis, exclusively upon inhibition of contractility (**Fig. 2F and Movies S2 and S3**).

**Figure 2:**
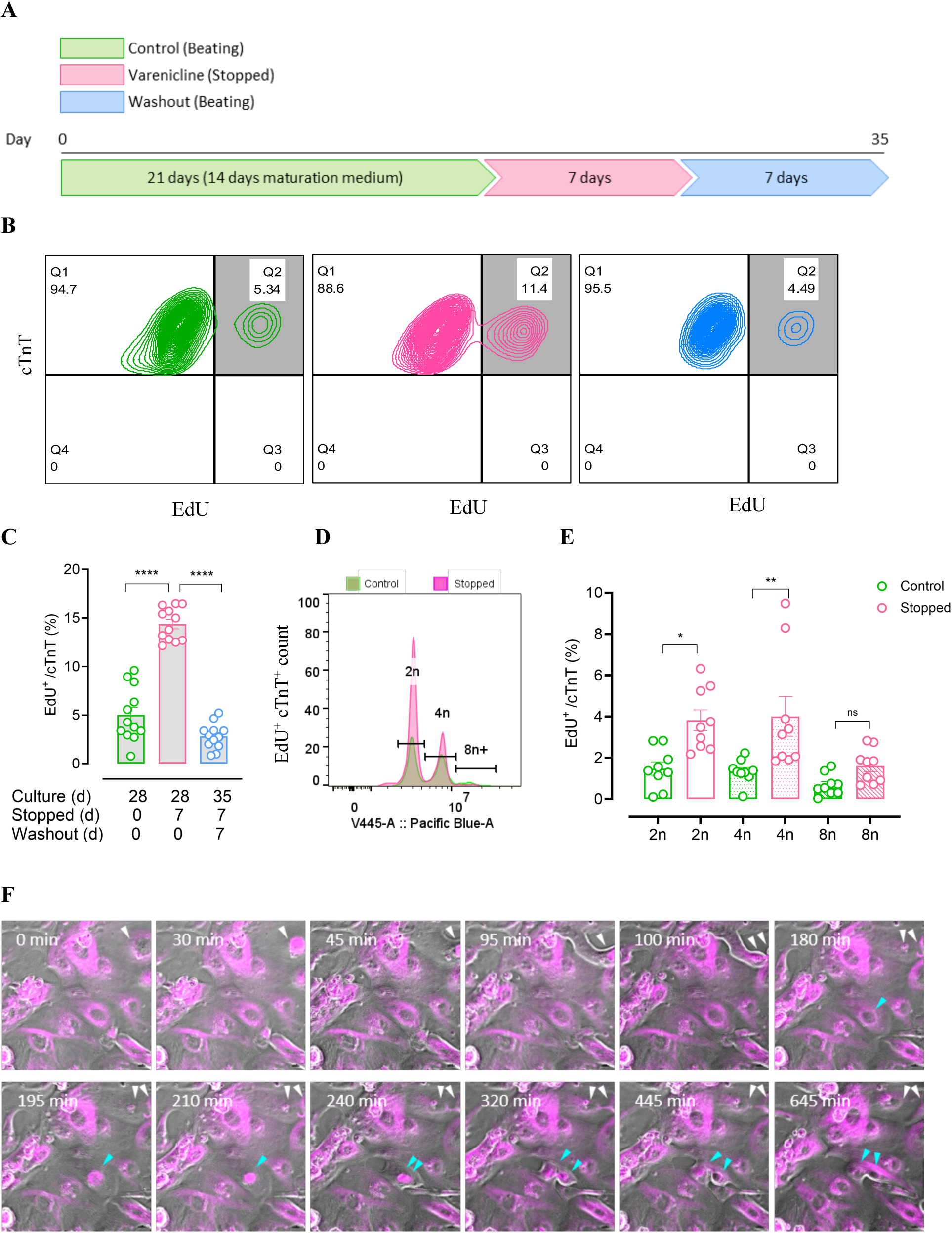
Cardiomyocyte proliferation in PSC-CMs was induced by contractility inhibition and reversed upon recovery. **(A**) Experimental timeline showing that cardiomyocytes were initially matured using a lipid-based maturation medium. EdU was applied for 24 hours prior to harvesting. (**B**) Flow cytometry analysis revealed increased cell cycle activity in PSC-cardiomyocytes following contractility suppression. Representative flow cytometry plots of EdU incorporation are shown before contractility inhibition, during inhibition, and after recovery. Only TnT-positive cells were analyzed. (**C)** Quantification of flow cytometry results (n = 12 biological replicates from 3 independent experiments). (**D**) Histogram of Hoechst DNA content in EdU⁺ cardiomyocytes under beating and contractility-inhibited conditions. (**E**) Quantification of Hoechst intensity in EdU⁺ cardiomyocytes from both conditions. (**F**) Time-lapse imaging captured cytokinesis in cardiomyocytes during contractility suppression. Cells were stained with SiR-tubulin and imaged over several days. Arrowheads indicate proliferating cardiomyocytes. Statistical analysis was performed using ANOVA. Significance levels are indicated as p <0.05 (*), p < 0.001 (**) and p < 0.00001 (****).

### Contractility controlled cardiomyocyte cell cycle activity in a 3D environment

To study whether contractility inhibition also stimulated proliferation in a more physiologically relevant context, we assessed cell cycle activity in EHTs under auxotonic load. Cardiomyocytes matured within the three-dimensional mechanically active environment, as indicated by cell alignment along the force vectors and time-dependent decline in cell cycle activity, measured by Ki67 expression (5.8% ± 0.9 at day 14 vs. 1.2% ± 0.3 at day 88; **Fig. S2 A**). Contractility was inhibited after a 21-day maturation period for durations of 1, 7, or 21 days, followed by re-initiation through agonist washout for 7 days, even after several weeks of continuous suppression **(Fig. 3A and Fig. S2 B-D)**. To investigate cardiomyocyte biology at a more advanced maturation stage, the culture period was extended up to 85 days. Inhibition of contractility was fully reversible after 7-day washout (**Fig. 3C**). Inhibition of contractility at the intermediate maturation stage resulted in increased cell cycle activity as shown by elevated Ki67 and phospho-histone H3 (PH3) expression (Ki67^+^ nuclei: 5.4% ± 1.57 in beating control EHTs vs. 7.5% ± 1.62 in stopped EHTs, n=3/N=3; PH3^+^ nuclei 0.13% ± 0.03 in beating control EHTs vs. 0.25% ± 0.1 in stopped EHTs, n=3/N=3). Utilizing the reversibility of the chemogenetic system, we studied cell cycle activity 7 days after contractility re-initiation, revealing a marked reduction in cell cycle activity (Ki67: 3.2% ± 2.2 nuclei, n=3/N=3, PH3: 0.12% ± 0.04 nuclei, n=3/N=3; **Fig. 3D-G**). Notably, even a single day of contractility inhibition was sufficient to increase cell cycle activity and extending the inhibition period up to 21 days did not further enhance this effect (**Fig. S2**). We next extended our analysis to long-term cultured EHTs, maintained for two months. At this mature stage, baseline cell cycle activity was minimal (0.9% ± 1 Ki67^+^ nuclei and 0.02% ± 0.02 PH3 nuclei, n=3/N=3). Nonetheless, contractility inhibition significantly increased cell cycle activity (Ki67: 4.5% ± 2.7 positive nuclei, PH3: 0.3% ± 0.14 nuclei, n=3/N=3), which again declined following re-initiation of beating (Ki67: 0.8% ± 1 and PH3: 0.05% ± 0.04 nuclei, n=3/N=3; **Fig. 3D, E, H, and I)**.

**Figure 3:**
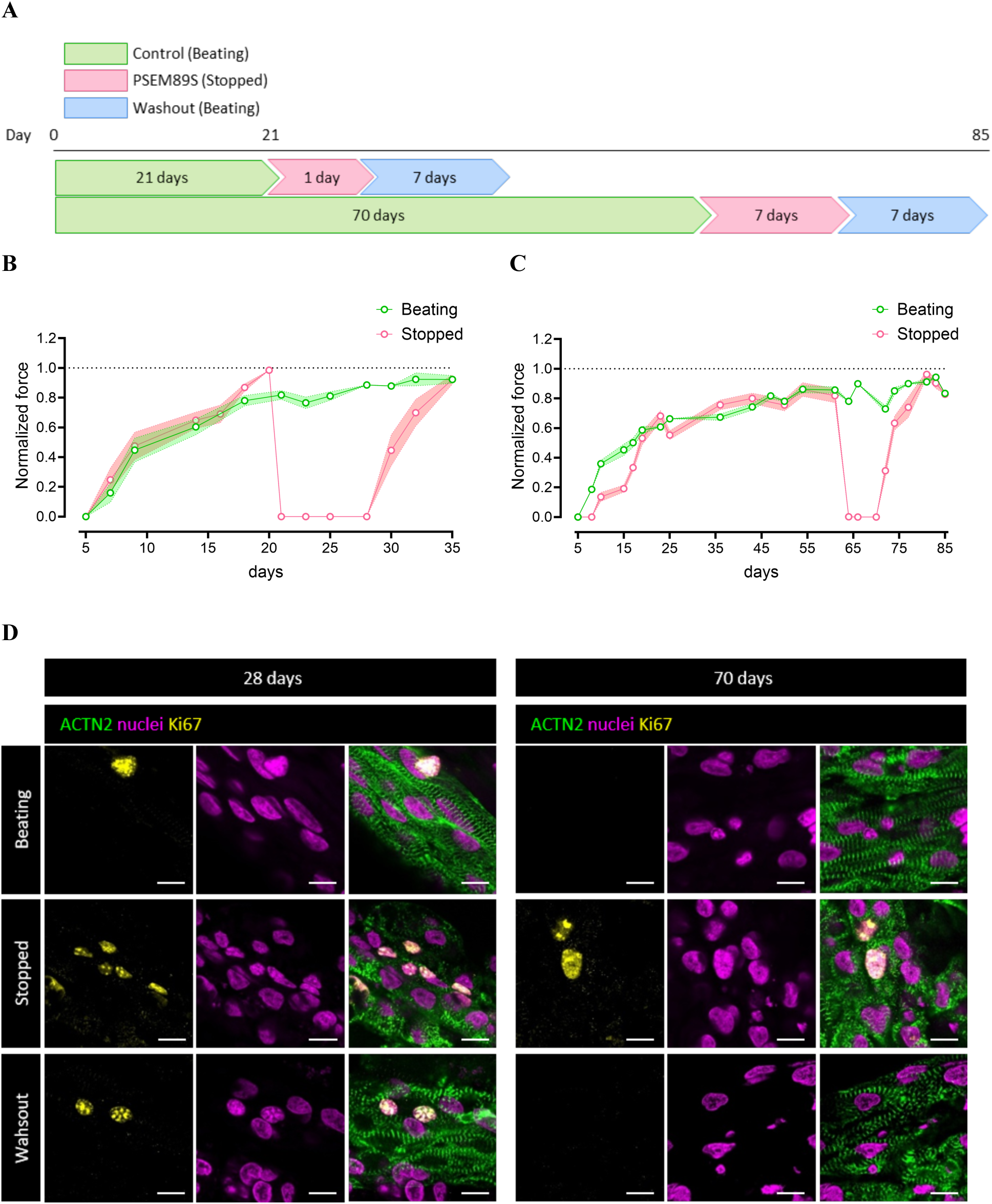
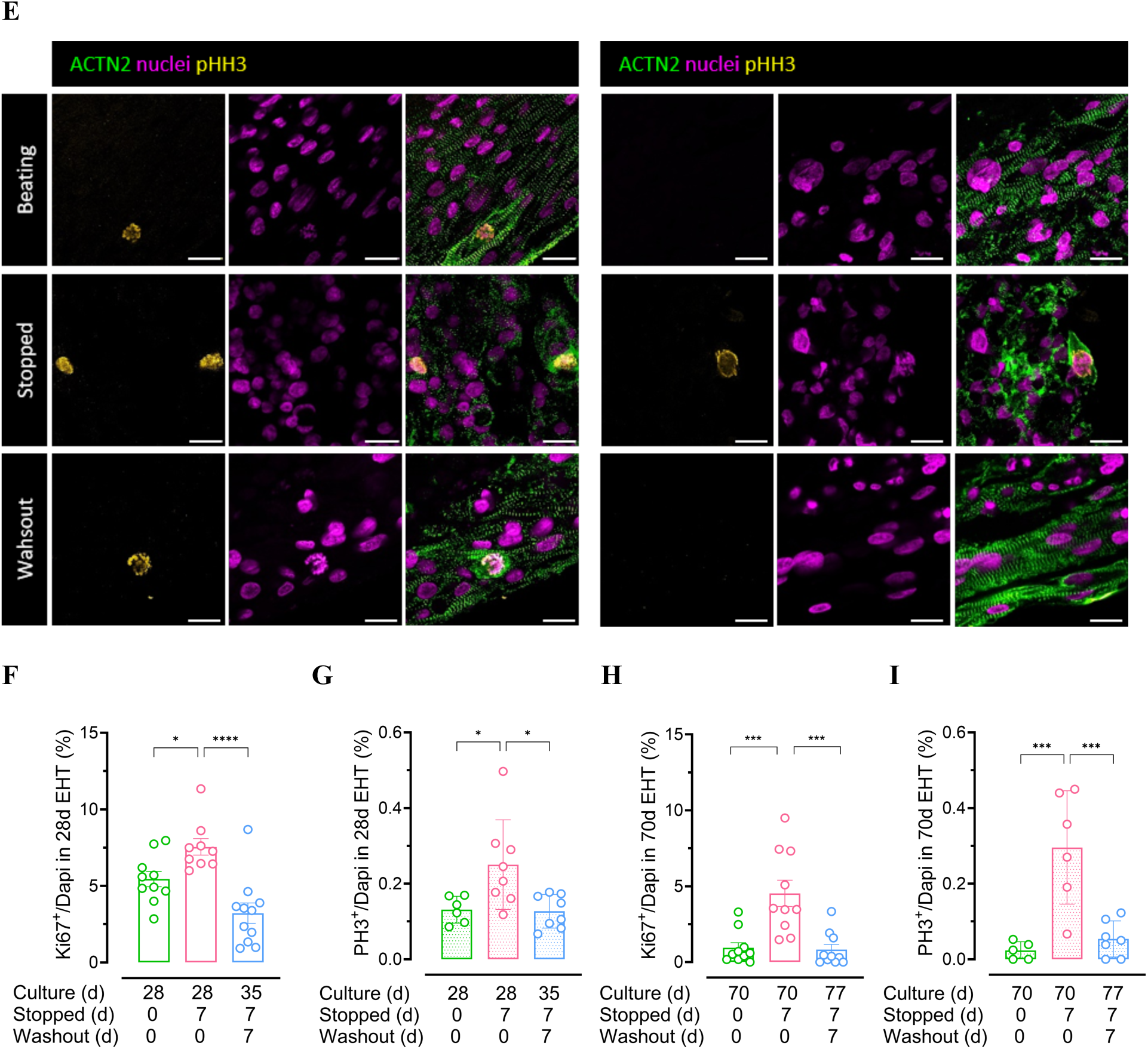
Cell Cycle Activity in human EHT is reversibly controlled by contractility. **(A**) Experimental timeline illustrating contractility inhibition applied for 7 days in EHTs cultured for 21 and 63 days, with an additional 7-day inhibition in matured EHTs. A 7-day washout period followed to assess recovery. **(B-C**) Contractile force recordings demonstrating inhibition of contractility and fully recovery after agonist washout in EHTs after a culture time of 28 days and 70 days respectively. **(D-I)** Contractility inhibition increased cell cycle activity, as evidenced by elevated expression of Ki67 **(D)** and PH3 (**E**). Cell cycle activity subsequently declined upon contractility recovery. (**F–I**) Quantification of Ki67 and PH3 positive nuclei. Each data point represents one EHT. For Ki67, 5 images per EHT were analyzed; for PH3, tiles imaging was performed and EHT section analyzed. Sample sizes ranged from 3 to 12 EHTs derived from 1 to 3 independent batches. PSAM^4^-GlyR cardiomyocytes were used for these experiments. Statistical analysis was performed using ANOVA. Significance levels are indicated as p < 0.05 (*), p < 0.0001 (***) and p < 0.00001 (****).

### Contractility inhibition stimulated cell cycle activity in adult human cardiomyocytes

Human myocardial slices from explanted end-stage failing hearts were transduced with PSAM^4^-AAV6 to enable chemogenetic control of contractility (**Fig. 4A**). Varenicline (100 nM, 7 days) reduced contractile force by 1.2 ± 0.3 mN relative to controls (n = 4/N = 3; **Fig. 4B, E**). Transduction efficiency was modest (7.7% ± 0.7, n = 4/N = 3), permitting assessment of contractility inhibition in individual cardiomyocytes within intact, beating tissue (**Fig. C, D and Fig S3**). PH3⁺ cardiomyocyte nuclei increased in treated slices (8.2% ± 1.1, n = 4/N = 2; **Fig. 4F),** predominantly within transduced RFP⁺ cardiomyocytes (52.9% ± 16.3, n = 4/N = 2; **Fig. 4G**) but also among non-transduced cardiomyocytes, indicating that both intrinsic and tissue-level contractility regulate cell cycle entry (**Fig.4H**). Independent antibody validation confirmed the unexpectedly high PH3 signal, establishing that contractility inhibition stimulates cell cycle activity in adult human cardiomyocytes.

**Figure 4:**
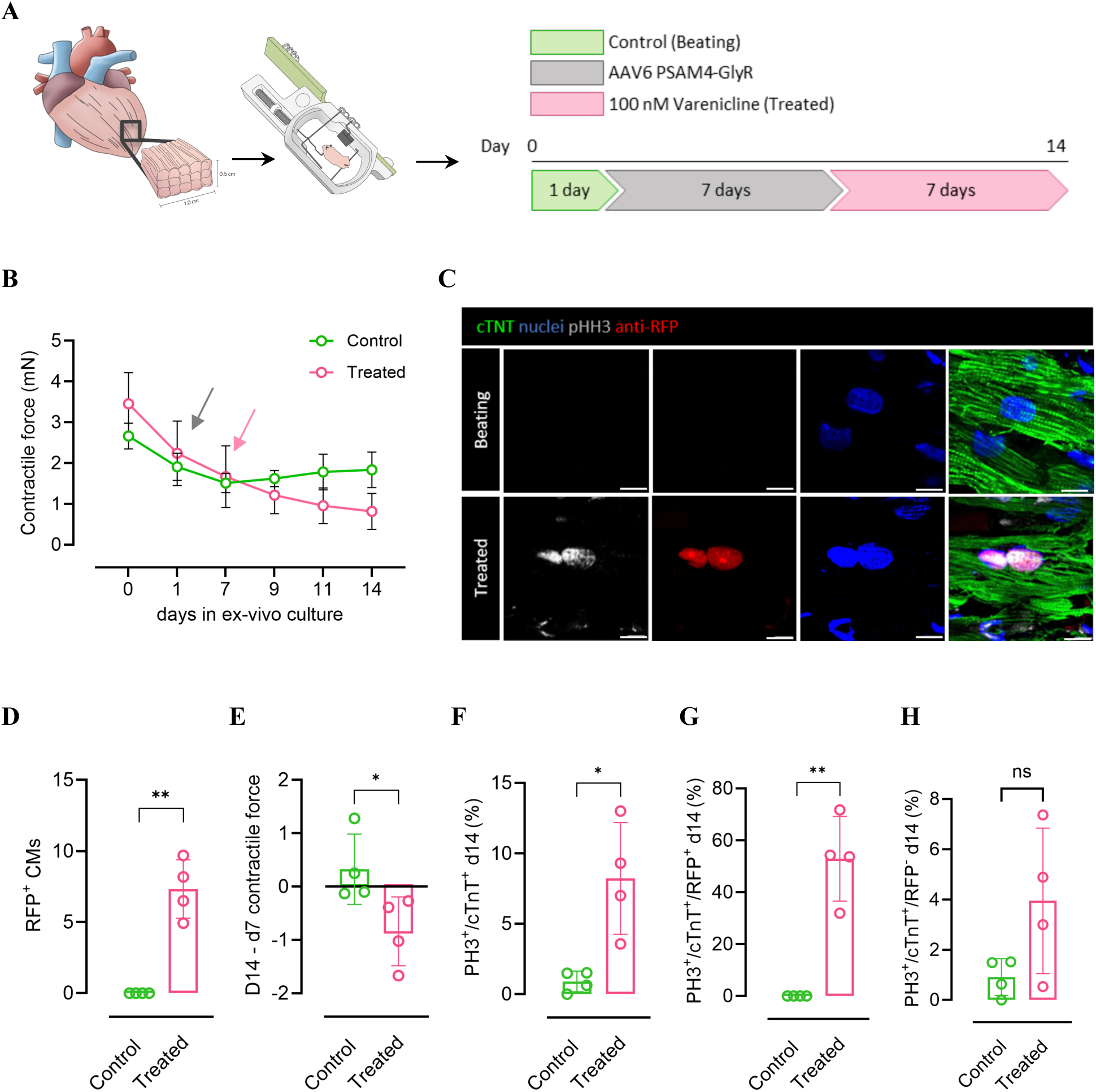
Contractility suppression induces proliferative activity in native cardiomyocytes. **(A)** Schematic of experimental design: human heart slices were transduced with AAV6-PSAM^4^-GlyR (Treated) and exposed to varenicline (100 nM) or left unmodified (Control). **(B)** Time course of contractile force over 14 days in culture showing progressive decline in treated slices compared with control, grey arrow represents onset of transduction and pink arrow represents onset of treatment. **(C)** Representative immunofluorescence of CTRL and treated slices stained for cTNT (green), nuclei (blue), PH3 (white), and RFP (red). **(D)** Percentage of RFP⁺ cardiomyocytes confirming AAV6 transduction. **(E)** Quantification of the change in contractile force after treatment. **(F-H)** Quantification of proliferative cardiomyocytes at day 14. **(F)** Percentage of PH3⁺cTNT⁺ cardiomyocytes. **(G)** PH3⁺cTNT⁺ nuclei restricted to RFP⁺ (transduced) cardiomyocytes. **(H)** PH3⁺cTNT⁺ nuclei in RFP⁻ (non-transduced) cardiomyocytes. Scale bars, 10 μm. Statistical analysis was performed using Students t-test. Significance levels are indicated as p < 0.05 (*), p <0.0.01 (**).

### Pharmacological contractility suppression increased cell cycle activity

We next investigated whether contractility inhibition via a depolarization-independent mechanism could also promote cardiomyocyte cell cycle activity. To this end, we employed mavacamten, a myosin ATPase inhibitor recently approved for the treatment of hypertrophic cardiomyopathy. Mavacamten stabilizes the myosin heavy chain in an energy-conserving state, termed the super relaxed state (SRX), thereby reducing contractile activity without affecting membrane potential (^32^). At a supratherapeutic concentration (10 µM), mavacamten stopped contractility; however, its onset and offset were considerably slower than the almost instantaneous contractile arrest observed with the chemogenetic approach, requiring approximately 1 hour to suppress beating and about 20 hours for recovery (**Fig. 5A**). At lower, therapeutically relevant concentrations (0.3 and 0.7 µM), mavacamten reduced contractile force in a concentration-dependent manner without fully abolishing beating (**Fig. 5B**). Consistent with findings from the chemogenetic approach, mavacamten-induced suppression of contractility led to sarcomere disorganization (**Fig. 5C, D**). This reduction in contractility was also associated with increased cell cycle activity (Ki67^+^ nuclei: 2.4% ± 0.3 in beating EHTs vs. 6.3% ± 0.2 in stopped EHTs, n=3/N=3; PH3^+^ nuclei: 0.18% ± 0.1 vs. 0.97% ± 0.2, n=3/N=2, respectively) (**Fig. 5E, F**). Fully recovered EHTs after mavacamten-washout led to cell cycle exit (Ki67+ nuclei: 1.5% ± 0.1, n=2) (**Fig. 5G**). Lower concentrations (0.3 and 0.7 µM) preserved beating while still reducing contractile force in a concentration-dependent fashion. Correspondingly, sarcomere disassembly and cell cycle activation also followed a concentration-dependent trend: cardiomyocytes treated with 0.3 µM retained sarcomere integrity and showed no significant increase in cell cycle activity, whereas 0.7 µM treatment led to sarcomere disorganization and enhanced cell cycle entry (**Fig. 5C-F**).

**Figure 5:**
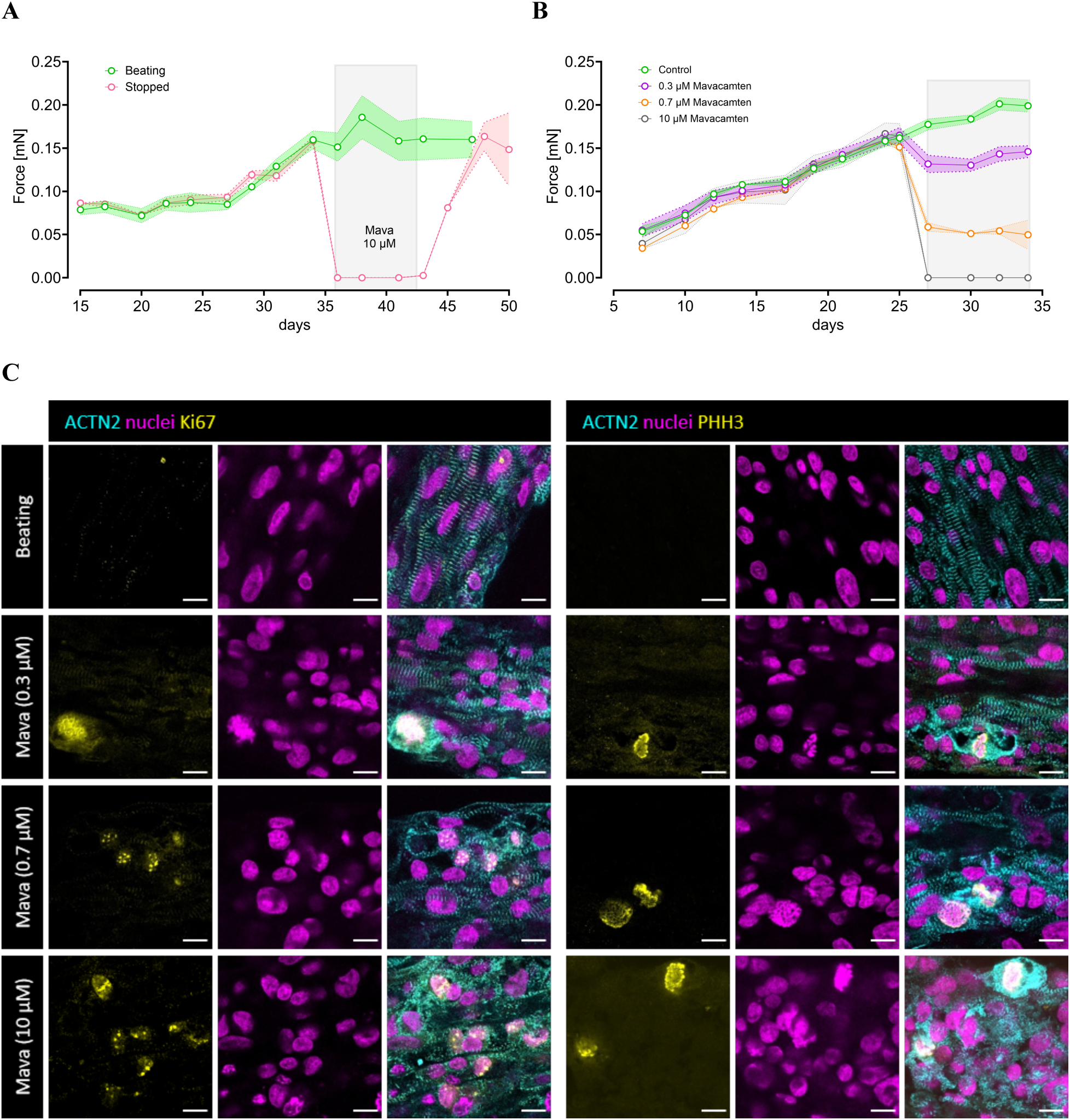
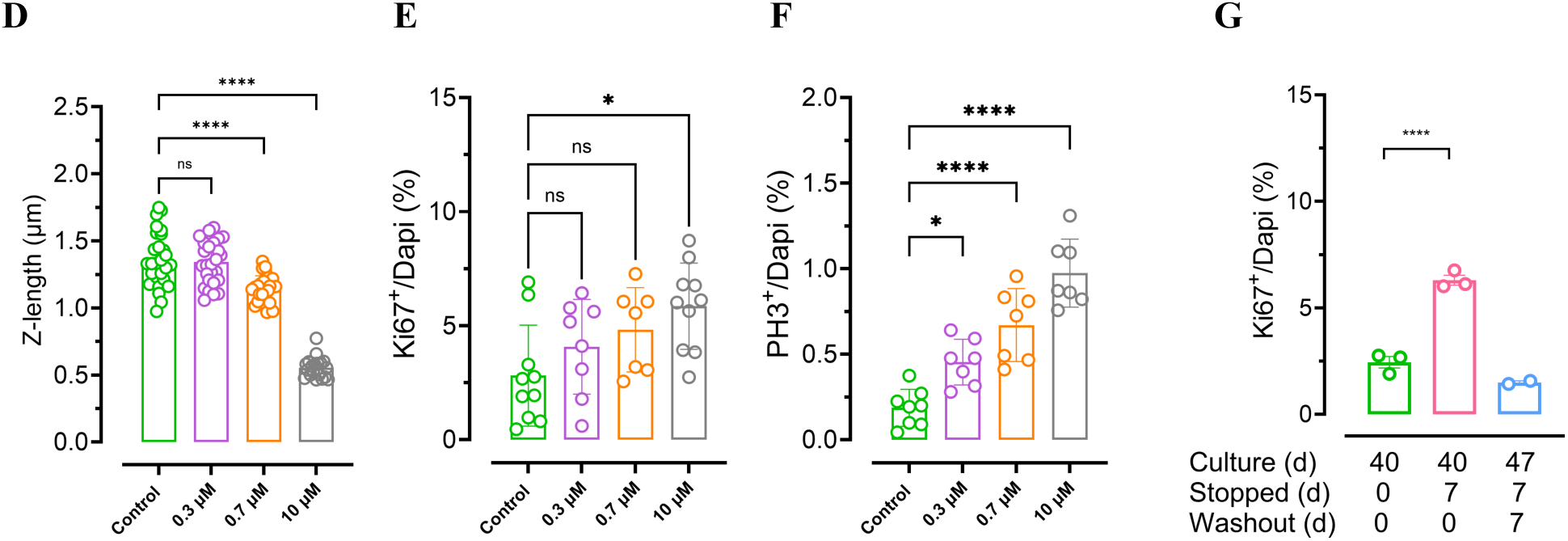
Mavacamten induced cell cycle activity in human EHT. **(A)** In matured EHTs, 10 µM mavacamten suppressed contractile force (day 36–43), which recovered after a 7-day washout **(B)** Time-course analysis of contractile force in EHTs treated with increasing concentrations of mavacamten (0.3, 0.7, 10 µM). Force decreased in a concentration-dependent manner. Shaded area indicates the treatment window. **(C)** Representative immunofluorescence images showing ACTN2 (cyan), Ki67 (yellow), and nuclei (magenta) in untreated (beating), mavacamten-treated (stopped) in different concentrations. (**D-F)** Quantification of sarcomere disassembly (Z-length, µm) and cell cycle activity (Ki67+ and PH3^+^ nuclei, %) at the different mavacamten concentrations (n=6-8). **(G)** Quantification of cell cycle activity (Ki67+ nuclei, %) before, during, and after mavacamten treatment. Ki67+ cells significantly increased during contractile inhibition and returned to baseline after recovery (n = 3 per group). Scale bar: 10 µm. Statistical analysis was performed using one-way ANOVA; significance levels: p < 0.05 (*), p < 0.00001 (****).

### Sarcomere disassembly preceded cardiomyocyte cell cycle re-entry

Live-cell imaging revealed not only an increase in cardiomyocyte proliferation but also provided mechanistic insights into the temporal relationship between contractility, sarcomere structure, and cell cycle activity. Contractility suppression occurred within seconds of agonist application, accompanied by rapid morphological and structural changes within the first few hours. However, initial signs of proliferation emerged only several hours later. To further investigate the kinetics of sarcomere disassembly and cell cycle re-entry, we performed histological analysis of PSAM^4^-GlyR cardiomyocytes at 1-, 6-, 12- and 24-hours following contractility inhibition. Sarcomere disassembly was evident as early as one hour after treatment, whereas increased cell cycle activity was only detectable after 24 hours (**Fig. 6A-D**), indicating that sarcomere disassembly preceded the cell cycle re-entry.

**Figure 6.**
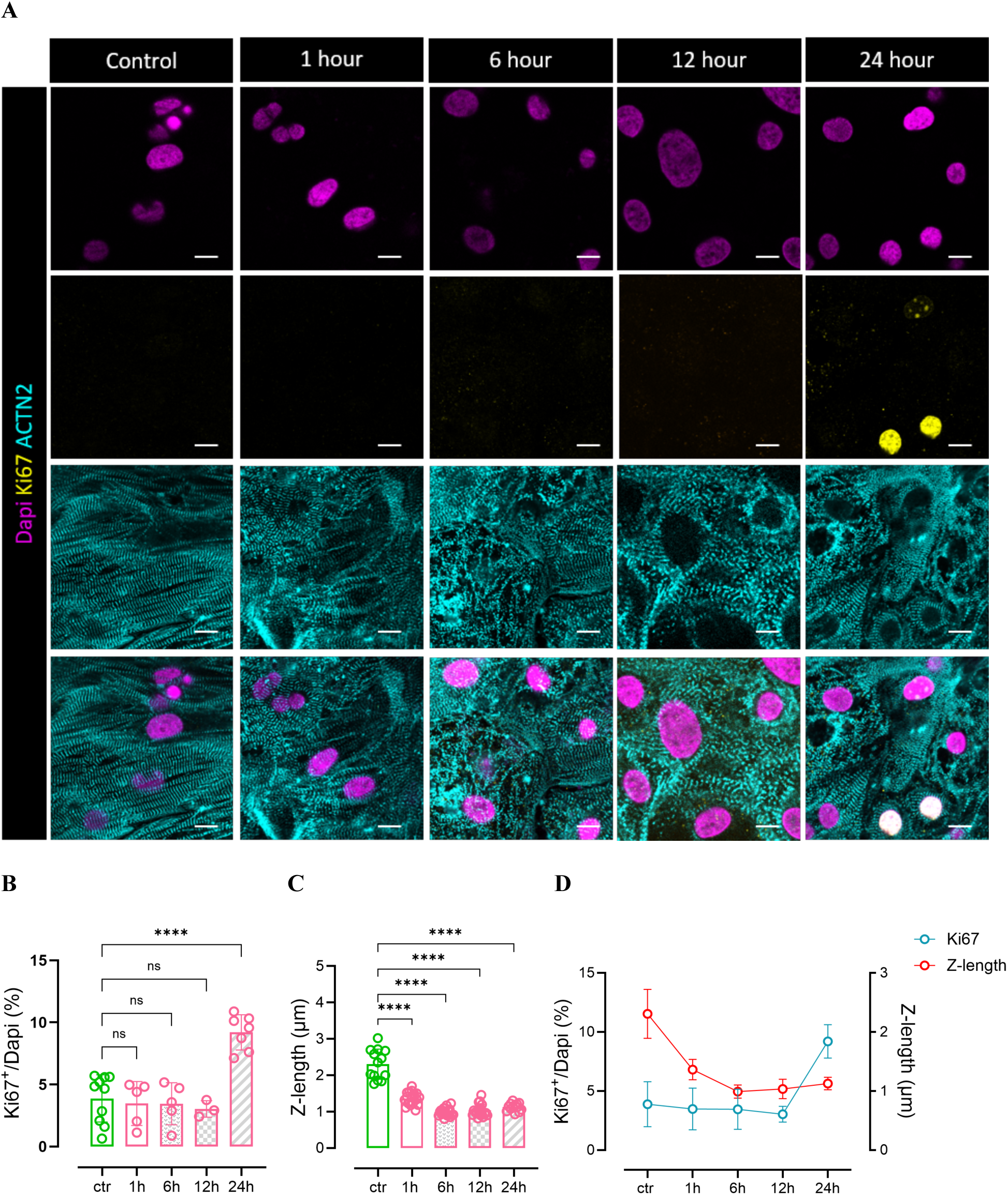
Sarcomere disassembly preceded cell cycle re-entry. **(A**) Time-course analysis of sarcomere structure and cell cycle activity in PSAM^4^-GlyR cardiomyocytes cultured in 2D. Sarcomere disassembly was detectable within the first hour after varenicline application, while increased cell cycle activity emerged only after 24 hours. **(B–D)** Quantification of sarcomere organization and cell cycle markers over time, confirming that structural disassembly precedes cell cycle re-entry. Scale bar = 20 µm. Statistical analysis was performed using one-way ANOVA; significance levels: p < 0.00001 (****).

### Transcriptional and proteomic profiling revealed SRF suppression and activation of E2F1/FoxM1 signalling in association with sarcomere disassembly and cardiomyocyte proliferation

To investigate molecular consequences of contractility inhibition, we performed both RNA sequencing and mass spectrometry-based proteomics on PSAM^4^-GlyR cardiomyocytes treated with varenicline (100 nM) for 7 days, compared to spontaneously beating controls and a 7-day recovery group (n=4 biological replicates per condition). Proteomic profiling via mass spectrometry supported the observation of sarcomere dysregulation showing significantly reduced expression of a broad set of sarcomeric proteins in non-beating CMs (**Fig. 7A**). Transcriptomic analysis revealed a regulation on the transcription level with a robust downregulation of genes encoding key sarcomeric components, including *MYH6*, *ACTN2*, and *TNNT2* (**Fig. 7B**). These alterations were consistent with histological observations. Transcriptional profiling further pointed to reduced activity of serum response factor (SRF) signalling, a well-established regulator of sarcomere structure and homeostasis. This finding was supported by our biochemical data showing increased levels of monomeric (G-) actin and reduced filamentous (F-) actin in non-beating CMs, as measured by Western blot (**Fig. 7C, D and Fig. S4**). This shift in actin dynamics is known to promote cytoplasmic retention of myocardin-related transcription factor (MRTF), thereby impairing SRF-mediated transcription. Pharmacological SRF inhibition with CCG-203971 (10 µM) disassembled sarcomeres but did not induce cardiomyocyte cell cycle activity (**Fig. S5**). Additionally, expression profiles shifted toward a more immature phenotype, as indicated by decreased *MYH7/MYH6*, *TNNI3/TNNI1*, *MYL3/MYL4* and *MYL2/MYL7* ratios (**Fig. 7E-H**). This dedifferentiation program also included marked downregulation of oxidative phosphorylation (OXPHOS)-related genes, reflecting a metabolic regression to a less mature state (**Fig. 7E**). In parallel, genes associated with cell cycle progression and DNA replication, including *CCNB1*, *CCNA2*, and *MKI67*, were significantly upregulated (**Fig. 7I**). Pathway analysis further implicated activation of *E2F1* and *FoxM1* signalling, both known drivers of cardiomyocyte cell cycle re-entry and proliferation. Gene ontology enrichment analysis supported these findings, highlighting pathways related to cytoskeletal remodeling, mitochondrial function, mitotic processes, and cell-cell interactions (**Fig. 7J-K**).

**Figure 7:**
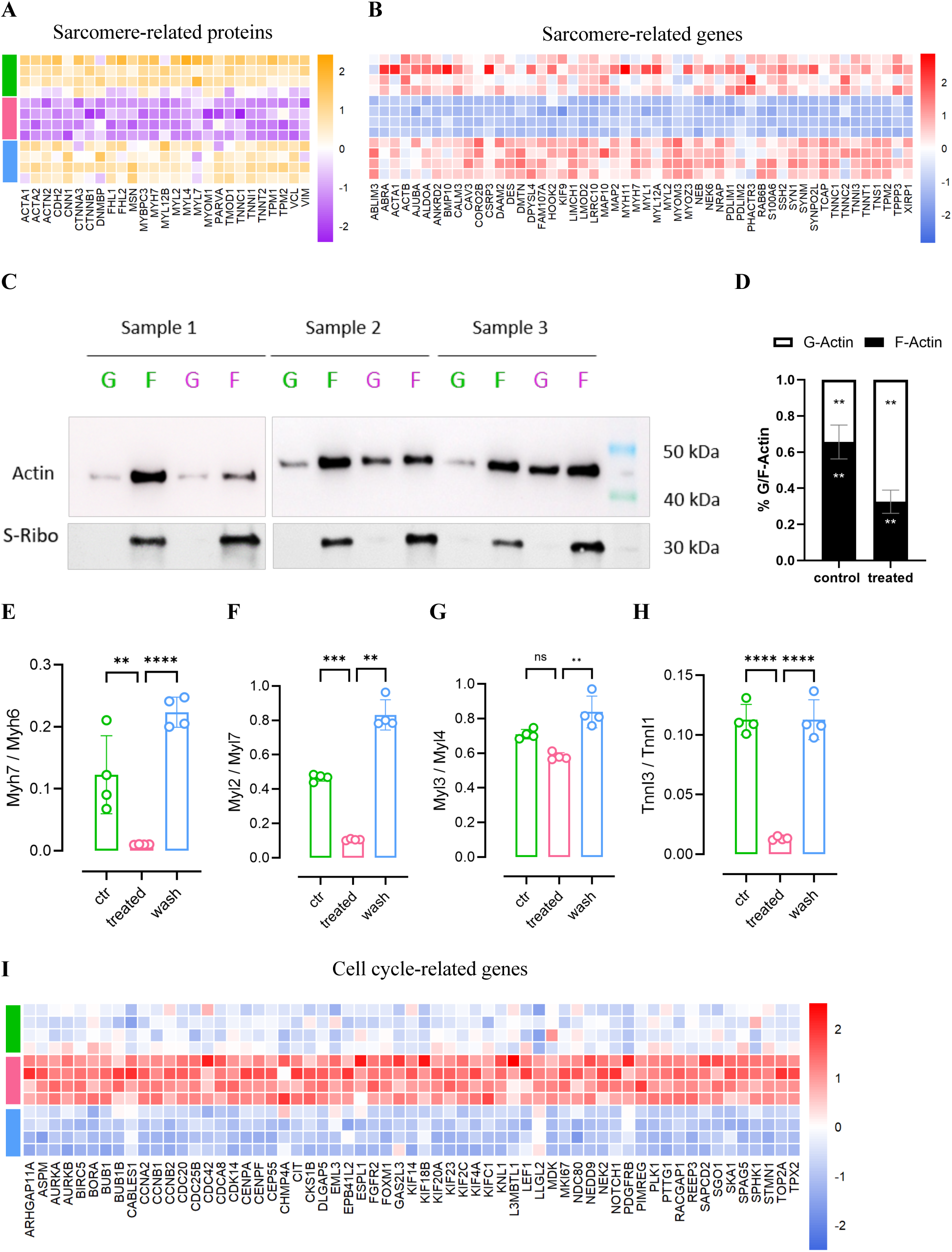
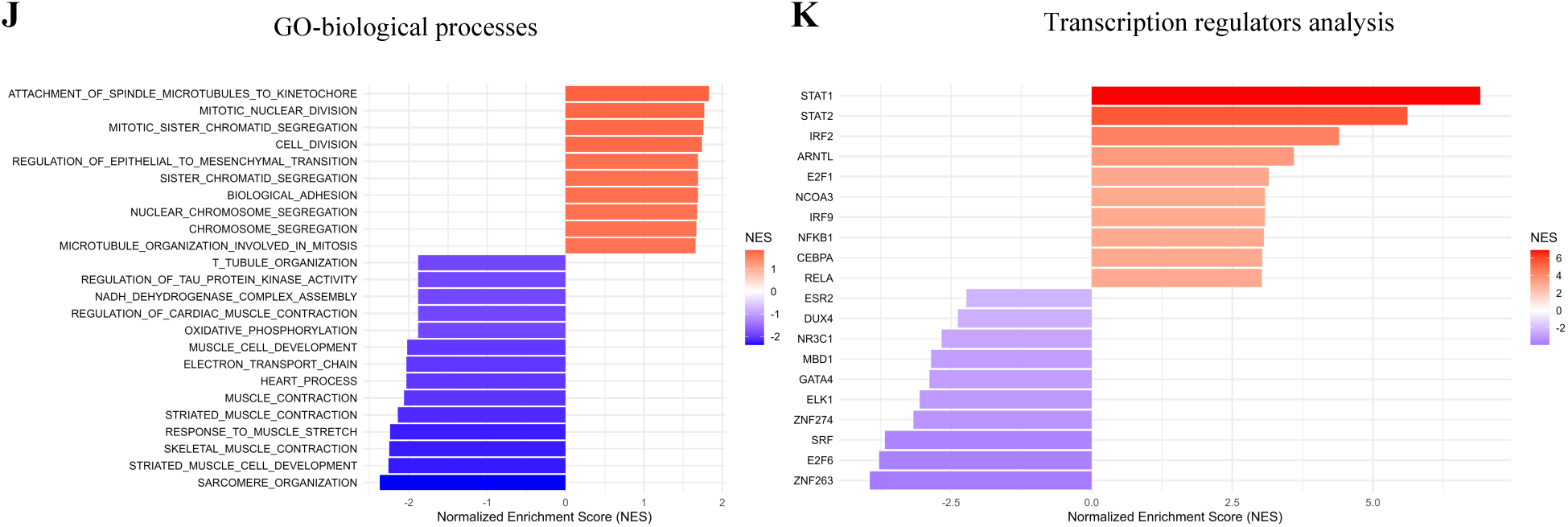
Multi-omics profiling reveals disrupted sarcomere homeostasis and increased cell cycle activity following contractility inhibition. **(A and B)** Proteomics and RNA sequencing analyses revealed disrupted sarcomere homeostasis, indicated by reduced abundance of sarcomeric proteins and decreased expression of sarcomeric genes. Comparisons were made across (beating) cardiomyocytes, non-beating (stopped) cardiomyocytes, and cardiomyocytes following contractility recovery (washout). (C-D) Western blot analysis demonstrated an increased G-actin to F-actin ratio during contractility inhibition. S6-Ribosomal protein was used as a housekeeping protein. (E-H) Isoform analysis showed a shift toward a more immature sarcomere gene expression profile upon contractility inhibition. (I) RNA sequencing further supported enhanced cell cycle activity, as indicated by upregulation of a broad range of cell cycle–related genes in contractility-inhibited cardiomyocytes. (J) Gene ontology representing up-and-down regulated biological processes when stopped CM are compared to beating control. (K) Transcription regulators analysis representing the transcription regulators expected to influence stopped vs beating CM.

## Discussion

The adult mammalian heart exhibits a very limited capacity for cardiomyocyte renewal, with turnover rates estimated at approximately 1% per year (^33–35^). While heart growth during embryonic development is driven by cardiomyocyte proliferation, these cells exit the cell cycle shortly after birth. Sedmera and Thompson noted that “what causes myocytes to divide is interesting, but what makes them stop dividing is fascinating” (^36^). This cell cycle exit coincides with profound physiological changes at birth as the neonatal circulation rapidly adapts to the increased hemodynamic demands of postnatal life: Left ventricular afterload increases with the loss of placental circulation, and cardiac output rises sharply (^37–39^). These hemodynamic changes impose a substantial increase in cardiac workload, temporally aligning with cardiomyocyte cell cycle exit (^40^), suggesting that increased mechanical load may contribute to the stop of cardiomyocyte proliferation.

Several indirect lines of evidence have supported a link between contractility and cell cycle regulation. Sarcomere disassembly, and a corresponding transient decline in contractile function, is a consistent feature of both natural and experimentally induced cardiomyocyte proliferation, suggesting a functional trade-off between force generation and proliferative potential (^41^). Morphological studies have documented the ordered disassembly and reassembly of sarcomeres during proliferation (^42–44^). Despite these observations, establishing a causal relationship between contractile activity and cell cycle arrest has remained elusive, largely due to the continuous demand for cardiac contraction and the technical challenges of selectively modulating contractility in a physiologically relevant context.

Here, we provide the direct experimental evidence that active contractility suppresses cardiomyocyte cell cycle activity. Using chemogenetic tools to precisely inhibit contractile function in hiPSC-derived cardiomyocytes, 3D-EHTs, and adult human cardiac slices, we establish a mechanistic link between intrinsic contractile activity and cell cycle regulation. Across all three models, inhibition of contractility led to a marked increase in cell cycle activity. Critically, the reversible nature of the chemogenetic system enabled us to demonstrate that restoring contractility prompted a return to cell cycle arrest, reinforcing the causal relationship between contractile function and cell cycle suppression. Two observations are particularly noteworthy: first, live-cell imaging demonstrated that contractility inhibition induced bona fide cardiomyocyte proliferation, including cytokinesis. Second, even terminally differentiated adult human cardiomyocytes within ex vivo cardiac slices responded to contractility inhibition with increased cell cycle activity. Notably, only a small subpopulation of cardiomyocytes had been transduced, while the surrounding tissue remained contractile. The transduced subpopulation exhibited an unexpectedly high increase (consistent with prior reports of pro-proliferative interventions in this system (^45^), accompanied by a concomitant rise in cell cycle activity among non-transduced cardiomyocytes. These findings indicate that intrinsic contractile activity is the primary regulator of cardiomyocyte proliferation, while reduced overall tissue-level contractility also provides an additional modulatory effect on cell cycle activity. The chemogenetic results were further corroborated by pharmacological inhibition using mavacamten, a selective myosin ATPase inhibitor, which similarly enhanced cell cycle activity. Together these data show that suppression of contractile activity, not modulation of electrical excitability, promoted cell cycle re-entry.

The structural changes observed, particularly sarcomere disassembly preceding cell cycle activation, point toward a mechanism in which contractile inhibition initiates sarcomere remodeling that in turn facilitates cell cycle reactivation. To explore the molecular underpinnings of this response, we employed a multi-omics approach. Proteomic analysis revealed alterations in RNA processing and chromatin remodeling pathways, consistent with changes in nuclear morphology, features known to be influenced by mechanical cues (^46^). These observations may reflect the involvement of nuclear mechanotransduction, as physical connections between the cytoskeleton and nuclear envelope provide a means for mechanical signals to influence nuclear architecture and function (^47^). Transcriptomic analysis further indicated that sarcomeric gene regulation occurred primarily at the transcriptional level, with pathway analysis implicating serum response factor (SRF) signalling. Pharmacological inhibition of SRF disrupted sarcomere structure, but did not increase cardiomyocyte cell cycle activity, consistent with findings from SRF knockout models (^48,49^). These results suggest that sarcomere disassembly alone, as also observed in titin-severing experiments (^50^), does not necessarily drive cardiomyocyte proliferation. Emerging evidence has identified adducin (particularly α- and γ-adducin) as potential mediators of sarcomere remodeling and cell cycle re-entry (^51^). Their role has been demonstrated in the regenerative neonatal heart, where the regenerative response was associated with increased adducin expression. This finding aligns with our proteomics data, which also showed elevated levels of α-adducin following contractility inhibition. Transcriptomic profiling revealed a shift toward a more immature gene expression signature, including a decreased *TNNI3/TNNI1* ratio, previously linked to enhanced proliferative capacity (^52^). Together, these findings suggest that reduced contractile demand regulated sarcomere homeostasis at the transcriptional level, not only driving general downregulation but also reactivating developmental gene programs linked to cardiomyocyte plasticity.

Our findings establish contractile activity as a physiological driver of cardiomyocyte cell cycle arrest and, by extension, a key contributor to the postnatal loss of regenerative capacity. By directly linking mechanical work to proliferation control, this study positions contractility as a potential therapeutic target for reactivating regenerative pathways in the adult heart. These insights support a model in which contractile demand triggers cardiomyocyte cell cycle exit, followed by the activation of transcriptional and structural programs that stabilize the non-proliferative state, programs that remain reversible upon removal of contractile load. Understanding and manipulating this sequence may be critical for advancing regenerative therapies for heart disease.

## Methods and materials

### Human iPSC culture and cardiac differentiation

Generation of the PSAM-GlyR and PSAM^4^-GlyR cell lines were recently described (^28,29^). In brief, the PSAM-GlyR or PSAM^4^-GlyR transgene was integrated in the AAVS1 locus of the UKEi001-A human iPSC line (hPSCreg: RRID:CVCL_A8PR). A master cell bank was generated at passage 37 and a working cell bank at passage 38. Karyotyping by Nanostring analysis was performed on the master cell bank and revealed a regular karyotype. Human iPSCs were maintained in FTDA medium on Geltrex-coated culture vessels (Gibco, A14133-02). For passaging, confluent hiPSC cultures were washed with PBS and dissociated using Accutase (Sigma-Aldrich, A6964-100mL) supplemented with 10 µM Y-27632 (Biorbyt, orb154626). SSEA3 analysis by FACS was performed prior to cardiac differentiation. To initiate cardiac differentiation, cells were collected by centrifugation at 200 × g for 2 minutes and resuspended in Stage 0 medium at a density of 3 × 10⁴ cells/mL. The cell suspension was transferred into spinner flasks (Integra, 182 051) placed on a magnetic stirrer (Variomag/Cimarec Biosystem Direct, Thermo Fisher Scientific, cat. no. 70101) within a hypoxia-controlled incubator and cultured for 24 hours at 40 rpm to facilitate embryoid body (EB) formation. On day 1, EBs were transferred to T175 suspension culture flasks at a density of 200– 250 µL EBs per 46 mL of Stage 1 medium and cultured for three days. On day 4, Stage 1 medium was replaced with FDM, and cultures were transferred to a normoxic incubator (21% O₂, 5% CO₂). On day 7, FDM was replaced with Stage II medium, with half-medium changes performed daily until day 11. Spontaneous contractions typically began around day 8. EB maturation was carried out in RDM medium until day 18. For dissociation, EBs were incubated in Collagenase II solution (200 U/mL in HEPES) supplemented with 30 µM N-benzyl-p-toluene sulfonamide (BTS, TCI, B3082-25G) and 10 µM Y-27632 for up to 4 hours at 37 °C. Following digestion, a blocking buffer was added, and cells were counted and processed for 2D culture or engineered heart tissue generation or cryopreserved.

### Cryopreservation

Cardiomyocytes were suspended in heat-inactivated FBS (Biochrom, S0615) + DMSO (Sigma-Aldrich, D4540) at 10:1 (∼3×10^7^ cells per cryovial). Cells were then transferred to -150 °C for long-term storage. Cryovials were transferred from -150 °C to a water bath of 37 °C until only a small ice clump remained. The cell suspension was then transferred to a bigger tube before drop-wise adding of 9 ml prewarmed RPMI medium supplanted with Penicillin/streptomycin 1% (v/v) and B27 10%. Cells were counted and used for 2D culture or EHT generation.

**Table S1.** Media composition for iPSC culture and cardiomyocyte differentiation.

| Medium | Composition |
| --- | --- |
| FTDA | DMEM/F-12 without glutamine (Life Technologies, 21331-046)<br>2 mM L-glutamine (Gibco, 25030-081)<br>5 mg/L Transferrin (Sigma-Aldrich, T8158)<br>6.6 µg/L Sodium selenite (Sigma-Aldrich, S9133-1MG)<br>1% (v/v) Human serum albumin (Biological Industries, 05-720-1B)<br>0.1% (v/v) Lipid mix (Sigma-Aldrich, L5146)<br>5 mg/L Human recombinant insulin (Sigma-Aldrich, I9278)<br>50 nM Dorsomorphin (Tocris, 3093)<br>2.5 µg/L Activin A (R&D Systems, 338-AC)<br>0.5 µg/L TGF β 1 (Peprotech, 100-21C)<br>30 µg/L bFGF (Peprotech, #100-18B) |
| Stage 0<br>(EB-formation) | FTDA<br>4 g/L Polyvinyl alcohol in 1x DPBS<br>10 µM Y-27632<br>30 µg/L bFGF |
| Stage 1<br>(EB-mesoderm induction) | RPMI 1640<br>4 g/L Polyvinyl alcohol (Sigma-Aldrich, P8136)<br>10 mM HEPES (pH 7.4) (Sigma-Aldrich, 9105.4)<br>0.05% (v/v) Human serum albumin<br>5 mg/L Transferrin<br>6.6 µg/L Sodium selenite<br>0.1% (v/v) Lipid mix<br>10 µM Y-27632<br>250 µM Phosphoascorbate (Sigma-Aldrich, 49752)<br>5 µg/L bFGF<br>3 µg/L Activin-A<br>10 µg/L BMP-4 (R&D Systems, 314-BP) |
| FDM<br>(EB cardiac specification) | RPMI 1640<br>0.5% (v/v) Penicillin/streptomycin (Gibco, 15140-122)<br>10 mM HEPES (pH 7.4)<br>0,5% (v/v) HSA<br>5 mg/L Transferrin |
|  | 6.6 µg/L Sodium selenite<br>0.1% (v/v) Lipid mix<br>1 µM Y-27632<br>250 µM Phosphoascorbate<br>1 µM XAV-939 (Tocris, 3748) |
| Stage II<br>(EB cardiac specification) | RPMI 1640<br>0.5% (v/v) Penicillin/streptomycin<br>10 mM HEPES (pH 7.4)<br>1 µM Y-27632<br>500 µM 1-Thioglycerol (Sigma-Aldrich, M6145)<br>2% (v/v) B27 with insulin (in-house production)<br>1 µM XAV-939 |
| RDM<br>(EB cardiac specification) | RPMI 1640<br>0.5% (v/v) Penicillin/streptomycin<br>10 mM HEPES (pH 7.4)<br>1 µM Y-27632<br>500 µM 1-Thioglycerol<br>2% (v/v) B27 with insulin |
| Blocking buffer | RPMI 1640<br>6 ml/L DNase II Type V (Sigma-Aldrich, D8764)<br>1% (v/v) Penicillin/streptomycin |

### Engineered heart tissue generation and functional characterization

EHTs were generated as previously described (^52^). PSAM-GlyR or PSAM^4^-GlyR cardiomyocytes were digested with collagenase II (Worthington, LS004176; 200 U/ml) in Ca^2+^-free HBSS (Gibco, 14175-053) with 1 mM HEPES (pH 7.4), 10 μM Y-27632, and 30 μM N-benzyl-p-toluene sulfonamide (TCI, B3082) for 3.5 hours at 37 °C (5% CO_2_, 21% O_2_). The dissociated cells were resuspended in Ca^2+^-containing DMEM with 1% penicillin/streptomycin. EHTs were generated in agarose casting molds with solid silicone racks (100 μL per EHT, 1×10^6^ cells). The culture medium was changed three times per week. Analysis of contractile force was performed by video-optical recording. The contraction peak analysis was performed during spontaneous beating (^16^).

### Cardiomyocytes 2D-monolayer culture

PSAM^4^-GlyR cardiomyocytes were cultured in 12 well plates with DMEM supplemented with 10% horse serum (Gibco, 26050-088 LOT: 1590399), 1% Penicillin/streptomycin and 10 µg/ml insulin. Media changes were performed every other day during the first week. For the following two weeks, a fatty acid-rich medium was used every 5 days to drive maturation (^1^). 24 hours prior to harvesting 10 µM EdU (ThermoFisher Scientific, C10338) was added to the culture medium.

### Flow cytometry

Single-cell suspensions of iPSC-derived cardiomyocytes were fixed in Histofix (Roth A146.3) for 20 min at 4 °C and stained in a buffer, containing 5% FBS, 0.5% Saponin (Sigma Aldrich, S7900), 0.05% sodium azide (Sigma Aldrich, S2002) in PBS. Antibodies are listed in Table S2. Samples were analyzed with NovoCyte Quanteon Flow Cytometer, the NovoExpress Software and BD FlowJo V10. For cell cycle studies EdU incorporation was assessed. After cell permeabilization, click-IT reaction with was performed as per manufacturer protocol using a mastermix containing Alexa-Fluor azide and CuSO_4_. Samples were counterstained with and APC-cTNT antibody and DAPI/Hoechst (1:1000) for 1 hour, washed, and resuspended in 250 μL PBS. Data acquisition was performed using the Agilent Quanteon cytometer.

**Table S2.** Antibodies for flow cytometry.

| Antigen | Supplier | Titer |
| --- | --- | --- |
| SSEA3 (AlexaFluor 647) | BD Pharmingen, Cat. 561145 | 1:5 |
| Isotype SSEA3 | BD Pharmingen, Cat. 557714 | 1:40 |
| Cardiac troponin T (APC) | Miltenyi Biotec, Cat. 130-120-403 | 1:100 |
| Isotype cardiac troponin T | Miltenyi Biotec, Cat. 130-120-709 | 1:100 |

### Immunohistology

EHTs were fixed in Histofix (Roth A146.3) overnight at 4 °C. Then sections, 70 µm thick, were obtained using a vibratome. Permeabilization was achieved with 1% BSA (Serva, 11930) and 0.3% Triton X-100 (Sigma Aldrich, 93422) for 90 minutes at room temperature. Staining solutions were prepared using primary antibodies (as outlined in Table S3) in permeabilization buffer. Secondary antibodies were used in a 1:500-dilution.

**Table S3.** Antibodies for immunohistology.

| Antigen | Titer | Supplier |
| --- | --- | --- |
| Ki67 | 1:1000 | ThermoFisher, 14-5698-82 |
| α-Actinin | 1:800 | Sigma, A7811 |
| Titin-9 | 1:100 | Myomedix |
| Phalloidin | 1:250 | Thermo, A22284 |
| PH3 | 1:1000 | Cell signaling, 9701L |
| DAPI | 1:1000 |  |

### Live cell imaging

PSAM^4^-GlyR cardiomyocytes were cultured in a 96-well plate with glass bottom (Greiner, 655090). Varenicline (100 nM) treatment was initiated 12 hours prior to imaging (Sigma Aldrich, PZ0004) HEPES (20 mM; Roth, 9105.4), and SiR-Tubulin (100 nM; Spirochrome, SC002) was added to the media. Time lapse images were acquired every 15 minutes using the Nikon Eclipse Ti2 microscope in the Cy5 channel at 37 °C (5% CO_2_, 21% O_2_).

### Western blot

PSAM^4^-GlyR cardiomyocytes were dissociated using TrypLE 10X (Gibco, A1217701) and processed following the manufacturer’s protocol for the G-Actin and F-Actin in vivo assay (Cytoskeleton, BK037). Cell dissociation conditions were optimized to preserve cytoskeletal integrity, ensuring minimal disruption of actin dynamics. G-Actin were separated from F-Actin using 100,000 G ultracentrifuge for an hour. Proteins were separated on 4-15% precast polyacrylamide mini-gels (Bio-Rad; 4561086) and transferred by electroblotting on nitrocellulose membranes. S6-Ribosomal protein was used as a housekeeping protein (Cell signaling #2217). Proteins were visualized with Clarity Western ECL substrate (Bio-Rad; 1705061), and signals were detected with the ChemiGenius² Bio Imaging System (Bio-Rad).

### Cytotoxicity analysis

The culture medium was collected after three days of culturing. LDH concentration was measured with the CyQUANT™ LDH Cytotoxicity Assay Kit (Thermofisher, C20301) according to the manufacturer’s protocol. EHT groups sections were stained using the Click-iT™ Plus **T**erminal deoxynucleotidyl transferase d**U**TP **N**ick **E**nd **L**abeling (TUNEL) kit (Thermofisher, C10619) as a confirmatory procedure. Sections were permeabilized with 1% BSA (Serva, 11930) and 0.3% Triton X-100 (Sigma Aldrich, 93422) for 90 minutes at room temperature followed by the staining according to the manufacturer’s protocol. For troponin I (cTnI) release analysis, the EHTs culture media was collected after 24 h of culturing and transferred for analysis on ice. High sensitivity cTnI was quantified with the ARCHITECT STAT highly sensitive Troponin | immunoassay (Abbott Diagnostics, USA, ARCHITECT i2000SR). Calibration was performed according to the manufacturer’s instructions. The analytical measuring interval was 1.9–50,000 pg/mL (limit of detection, LoD: 1.9 pg/mL).

### LC-MS/-based bottom-up proteomics

Samples were prepared as described before (^53^). Cardiomyocytes were washed with PBS before harvesting and snap freezing. Samples were resuspended in 100 mM triethylammonium bicarbonate (TEAB) containing 1% (w/v) sodium deoxycholate (SDC), heat-denatured at 95 °C for 5 minutes and sonicated using a probe sonicator. Protein concentrations of denatured samples were determined using the Pierce BCA Protein Assay Kit (Thermo Fisher), and samples were normalized to contain 10 µg of total protein. Digestion was carried out in a 96-well LoBind plate (Eppendorf, Hamburg, Germany) using semi-automated pipetting via an Andrew+ Pipetting Robot (Waters, Milford, USA). Disulfide bonds were reduced in 10 mM dithiothreitol at 56 °C for 30 minutes with shaking (800 rpm) and alkylated with 20 mM iodoacetamide for 30 minutes at 37 °C. A 1:1 mixture of carboxylate-modified magnetic E3 and E7 SpeedBeads (Cytiva Sera-Mag™, Marlborough, USA) in LC-MS grade water was added at a 10:1 bead-to-protein ratio following a modified SP3 workflow (^54^). Protein binding occurred at 50% acetonitrile (ACN) while shaking at 600 rpm for 18 minutes. Beads were washed twice with 80% ethanol and once with 100% ACN. Proteins were digested overnight at 37 °C in 100 mM ammonium bicarbonate (AmBiCa) using sequencing-grade trypsin (Promega) at an enzyme-to-protein ratio of 1:100, with shaking at 500 rpm. Trypsin was inactivated by adding trifluoroacetic acid (TFA) to a final concentration of 1% and shaking for 5 minutes at 500 rpm. The resulting peptide-containing supernatants were transferred to a fresh 96-well LoBind plate for LC-MS/MS analysis. Peptides were separated chromatographically using a nano-UHPLC system (Dionex Ultimate 3000, Thermo Fisher) equipped with a two-buffer system (Buffer A: 0.1% formic acid (FA) in water; Buffer B: 0.1% FA in ACN). Online desalting and pre-concentration were performed using a trap column (100 µm × 20 mm, 100 Å, 5 µm C18, Nano Viper, Thermo Fisher), followed by separation on a 25 cm reversed-phase analytical column (75 µm × 250 mm, 130 Å, 1.7 µm BEH C18, nanoEase, Waters). Peptides were eluted over 80 minutes with a linear gradient from 2% to 30% buffer B over 60 minutes. Mass spectrometric analysis was performed on a Q Exactive hybrid quadrupole-Orbitrap mass spectrometer (Thermo Fisher Scientific) with nano-electrospray ionization (nano-ESI) at 1,800 V in data-dependent acquisition (DDA) mode. Full MS1 scans were acquired at a resolution of 70,000 (m/z 200), with a scan range of m/z 400–1,200, a maximum injection time of 240 ms, and an AGC target of 1 × 10^6. The top 15 most intense precursor ions (intensity threshold: 5 × 10^3; charge states: +2 to +5) were selected for fragmentation using higher-energy collisional dissociation (HCD) at a normalized collision energy of 25% and isolated with a 2 m/z window. MS2 scans were acquired at a resolution of 17,500, with a starting m/z of 100, a maximum injection time of 50 ms, and an AGC target of 1×10^5^. Fragmented precursors were excluded from re-selection for 20 seconds. Raw LC-MS/MS data were analyzed using the Sequest algorithm Homo sapiens (v2023-11-08), and common contaminants. Carbamidomethylation of cysteine residues was set as a fixed modification, while oxidation of methionine, methionine loss, and N-terminal acetylation were considered variable modifications. Up to two missed tryptic cleavages were allowed. Only peptides ranging from 6 to 144 amino acids were considered. A strict false discovery rate (FDR < 0.01) was applied at both the peptide and protein levels. Quantitative analysis was conducted using the Minora algorithm in Proteome Discoverer. Protein abundance values were log2-transformed and normalized via column-median normalization.

### RNA sequencing

To uncover the transcriptional mechanisms underlying the observed increase in cardiomyocyte proliferation, we performed RNA sequencing on CM cultured in monolayer for 4 different conditions; control (28d), treated for 7 days (28d), wash for 7 days (35d) and older control (35d). Total RNA was extracted from snap-frozen CM samples and assessed for integrity using the Agilent TapeStation system. High-quality RNA (RIN > 8) was poly(A)-enriched and used for directional library preparation (NEBNext Ultra II, NEB), followed by quantification and equimolar pooling. Libraries were sequenced on the Illumina NextSeq2000 platform (single-end, 72 cycles). Sequence reads were processed with fastp (v0.23.2) to remove sequences originating from sequencing adapters and sequences of low quality using the program’s default parameters. Mismatched base pairs in overlapped regions with one base with high quality while the other is with ultra-low quality were corrected with the --correction option (^54^). Reads were then aligned to the human reference assembly (GRCh38.110) using STAR (v2.7.10a) (^55^). Differential expression was assessed using DESeq2 (^56^). A gene was considered significantly differentially expressed if the corresponding absolute fold change was not less than 1.5 and the false discovery rate (FDR) did not exceed a value of 0.05. Overrepresentation (ORA) and gene set enrichment analysis (GSEA) was performed with clusterProfiler (^57^) in combination with a selection of the Molecular Signatures Database (MSigDB) subsets C2 and H, i.e. Biocarta (^58^), KEGG (^59^), Reactome (^60^), WikiPathways (^61^) and the Hallmark gene sets. The Gene Ontology (GO) resource (^62^) was analysed separately. Transcription factor activity predictions are based on DoRothEA (v1.6.0) (^63^).

### Ex vivo human heart slice culture

For ex vivo heart slice cultivation human tissue was obtained from patients undergoing heart transplantation at the University Medical Center Munich, Germany. Explanted human heart tissue was placed in 2,3-butanedione 2-monoxime (Sigma-Aldrich) at 4 °C. Tissues were sectioned on vibratome (Leica Biosystems) to 1 cm × 2 cm × 300 µm. Slices were anchored in biomimetic cultivation chambers (MyoDishes; InVitroSys) with tissue adhesive (histoacryl; B. Braun) according to fibre direction and subjected to physiological preload of 1 mN and stimulation at 1 Hz, as previously described (^64,65^). The MyoDishes are anchored on a rockerplate in an incubator. After the slices adapted to the biomimetic ex vivo culture, they were transduced with an adeno-associated virus serotype 6 (AAV6) vector encoding a cardiac-specific cTNT promoter-driven PSAM^4^-GlyR construct (rssAAV6-TnT-nls-TagRFP-2A-PSAM^4^-GlyR), produced by the Vector Core Facility, UKE. Transduction was confirmed microscopically after 24 hours. Treatment with varenicline (100 nM) was initiated on day 7 and continued for 7 days. Contractile force was continuously measured, and data were imported (MyoDish Data file converter v1.1) and analysed by LabChart Reader software (V8.1.14, AD Instruments). For immunofluorescence analysis, fixed samples were embedded in gelatin and cryosectioned into 14-µm-thick sections. The sections were blocked and permeabilized for 2 hours at room temperature in a solution containing 0.1% Triton X-100, 0.05% Tween 20, and 10% fetal bovine serum (Biophyll). All reagents were from Sigma-Aldrich unless otherwise specified. Primary antibodies were diluted in 0.1% Triton X-100, and 10% fetal bovine (according to Table S4) and incubated overnight at 4°C. Following incubation, sections were washed with a 0.05% Tween 20 solution. Secondary antibodies were then applied and incubated for 2 hours at room temperature. After several washing steps, nuclei were stained with Hoechst 33258. Slides were mounted with Anti-Fade Fluorescence Mounting Medium (Abcam, AB104135) and sealed with a coverslip. Images were acquired using two separate systems: a Leica DMI8 Thunder microscope and a Leica TCS SP8 confocal microscope. Image acquisition was performed using the LAS X software (Leica Microsystems). Image analysis was conducted with Fiji software.

**Table S4.** Antibodies for immunofluorescence in ex-vivo adult heart slices.

| Target | Host | Reference | Titer |
| --- | --- | --- | --- |
| Anti-Mouse IgG Alexa Fluor 488 | Goat | Invitrogen A11001 | 1:500 |
| Anti-Mouse IgG Alexa Fluor Plus 647 | Goat | Invitrogen A32787 | 1:500 |
| Anti-Rabbit IgG Alexa Fluor 647 | Goat | Invitrogen A31573 | 1:500 |
| Cardiac Troponin T | Mouse | Invitrogen MA5-12960 | 1:300 |
| Hoechst 33258 |  | Sigma-Aldrich 94403 | 1:500 |
| Phospho-Histone H3 Alexa Fluor 647 | Rabbit | Cell Signaling 9716S | 1:100 |
| RFP Booster ATTO594 | Alpaca | Proteintech rba594 | 1:100 |

### Quantification and statistical analysis

GraphPad Prism software version 9 (GraphPad Software, San Diego, CA, USA).

## Acknowledgments

We would like to acknowledge Thomas Schulze, Birgit Klampe, Elisabeth Kramer, Grit Höppner, Simona Parretta, and Moritz Meyer-Jens for technical assistance. We would like to appreciate the contribution of Dr. Sandra Laufer during reprogramming the UKEi001-A line. Flow cytometry was conducted in the FACS Core Facility, UKE. Image acquisition was conducted at the UKE and the CNIC Dynamic Imaging and Microscopy Unit.

## Funding

This work was supported by:

A Translational Research Grant from the German Centre for Cardiovascular Research (DZHK; 81X2710153 to TE)
The European Research Council (ERC-AG IndivuHeart to TE)
The European Union’s Horizon 2020 research and innovation program (874764 to TE)
The German Research Foundation (DFG; WE5620/3-1 to FW)
The European Uniońs Horizon2020 FetOpen RIA (964800 to FW)
Jung Career Advancement Award (to CMP)

CNIC is supported by:

The Instituto de Salud Carlos III (ISCIII)
The Ministerio de Ciencia, 437 Innovación y Universidades (MCIU, MICIU/ AEI/ 10.13039/ 501100011033)
The Pro CNIC 438 Foundation and a Severo Ochoa Center of Excellence (grant CEX2020-001041-S funded by 439 MCIU).
MALU is supported by grant FJC2021 047055-I from MCIU.
YR was supported by the La Caixa Foundation with a predoctoral fellowship.

## Author contributions

Conceptualization: NS and FW.

Funding acquisition: TE and FW.

Investigation: NS, RS, CM, TS, MN, AC, CM, BS, PN, AW, MZC, YR, LZG, JR, AGE, MAGR, RDM, EB, CMP.

Project administration: TE, FW

Resources: TE, FW.

Supervision: FW.

Visualization: NS, RS, FW.

Writing – original draft: NS, FW.

Writing – review & editing: , CMP, TE.

## Competing interests

T.E. and F.W. participate in a structured partnership between Evotec AG and the University Medical Center Hamburg-Eppendorf (UKE) on the development of a cell-based therapy for heart failure patients. A.D is a co-founder and shareholder in InVitroSys GmbH.

## Data and material availability

Requests for resources and reagents should be directed to and will be fulfilled by the lead contact, Florian Weinberger. The PSAM-GlyR and the PSAM^4^-GlyR hiPSC lines will be made available on request with a completed Materials Transfer Agreement.

## Supplementary Materials

Materials and Methods

Figs. S1 to S5

Tables S1 to S4

References (*53*–*65*)

Movies S1 to S3

Data S1 to S2

---

**Fig. S1.**
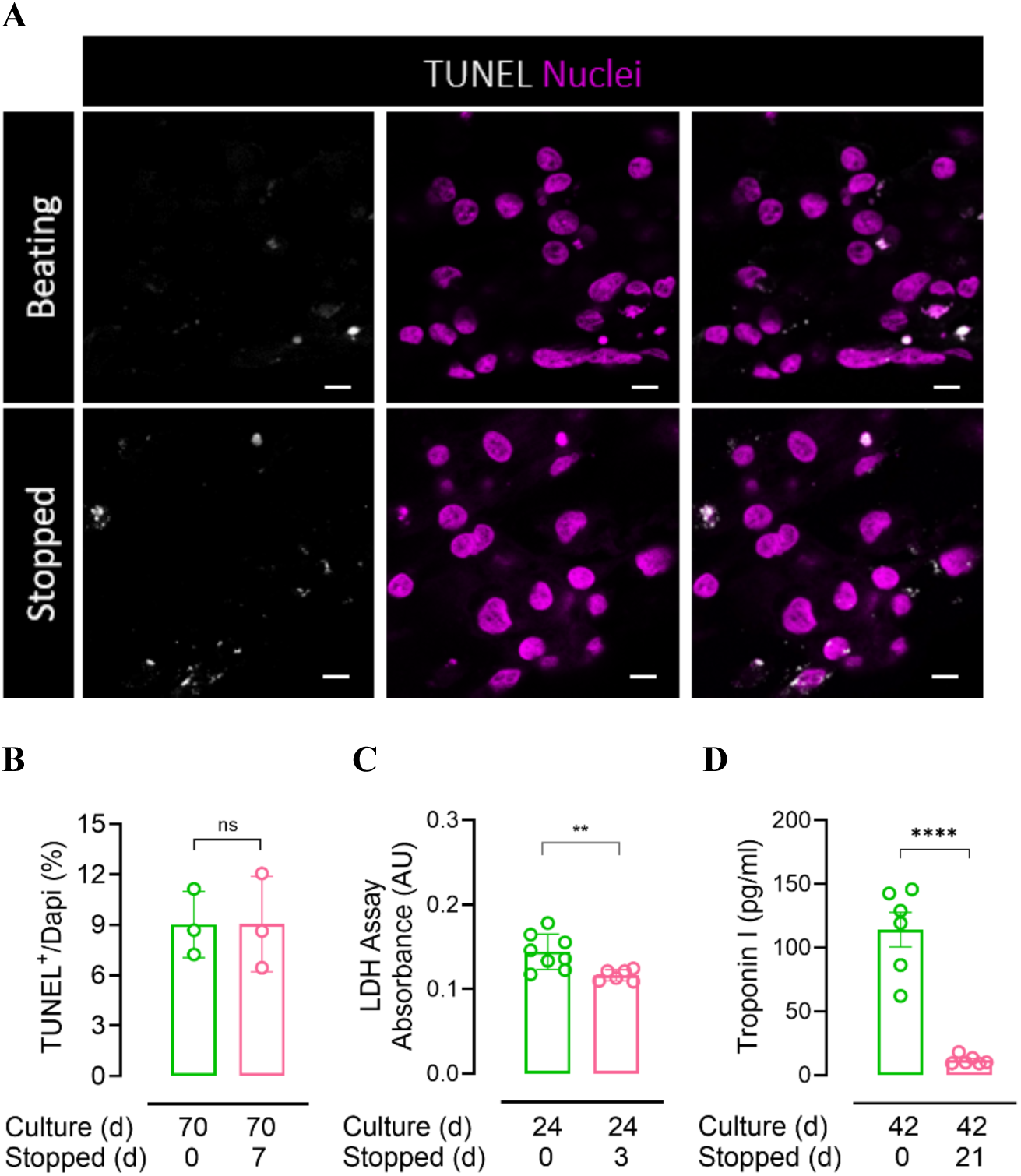
Contractility inhibition does not induce cell death in engineered heart tissue. **(A)** TUNEL staining images show no increase in TUNEL-positive nuclei in non-beating cardiomyocytes compared to beating controls. Nuclei are shown in magenta; TUNEL staining is shown in white. Scale bar = 20 µm. (**B**) Quantification of TUNEL staining confirms no increase in DNA fragmentation after 7 days of contractility inhibition (n = 3 biological replicates, each 5 images). (**C)** LDH release was slightly but significantly reduced after 3 days of contractility inhibition (n = 6 biological replicates). (**D)** Troponin I concentration in the culture medium were markedly decreased following 21 days of contractility suppression (n = 6 biological replicates). Statistical analysis was performed using unpaired t-tests. Significance levels: p < 0.05 (*), p < 0.0001 (***).

**Fig. S2:**
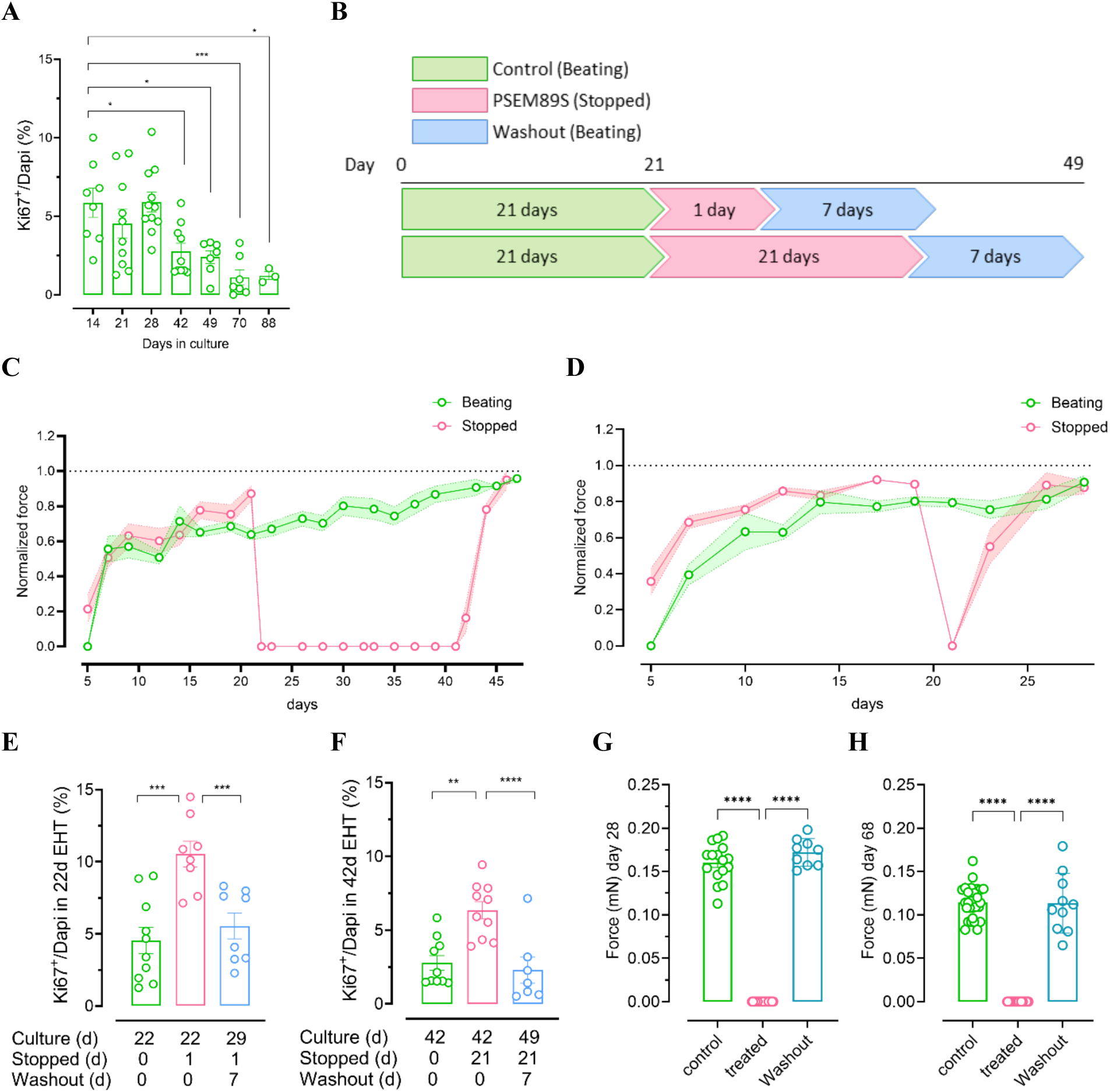
Cell Cycle Activity in human EHT is reversibly controlled by contractility. **(A)** Cultivation in a three-dimensional environment matured cardiomyocytes as indicated by a decrease in cell cycle activity measured by Ki67 expression. (**B)** Experimental timeline illustrating contractility inhibition applied for 1 and 21 days in EHTs cultured for 21 days. A 7-day washout period followed to assess recovery. Transcriptome analysis of sarcomere-related genes indicating down-regulation during contractility inhibition compared to beating CM (control or recovered). (**C-D)** Contractile force recordings demonstrating inhibition of contractility and fully recovery after agonist washout in EHTs after a culture time of 21 days and a suppression of contractility for 1 and 21 days respectively. (**E-F)** Quantification of cell cycle activity in PSAM^4^-GlyR EHTs stopped for 1 or 21 days respectively. Cell cycle activity was assessed by Ki67 expression (n=7-10 EHTs per group from 3 different batches, 5 images per EHT were assessed). (**G-H**) Absolute force measurements at day 28 and day 68 respectively. Statistical analysis was performed using one-way ANOVA. Significance levels: p < 0.001 (**), p < 0.0001 (***), p < 0.00001 (****).

**Fig. S3.**
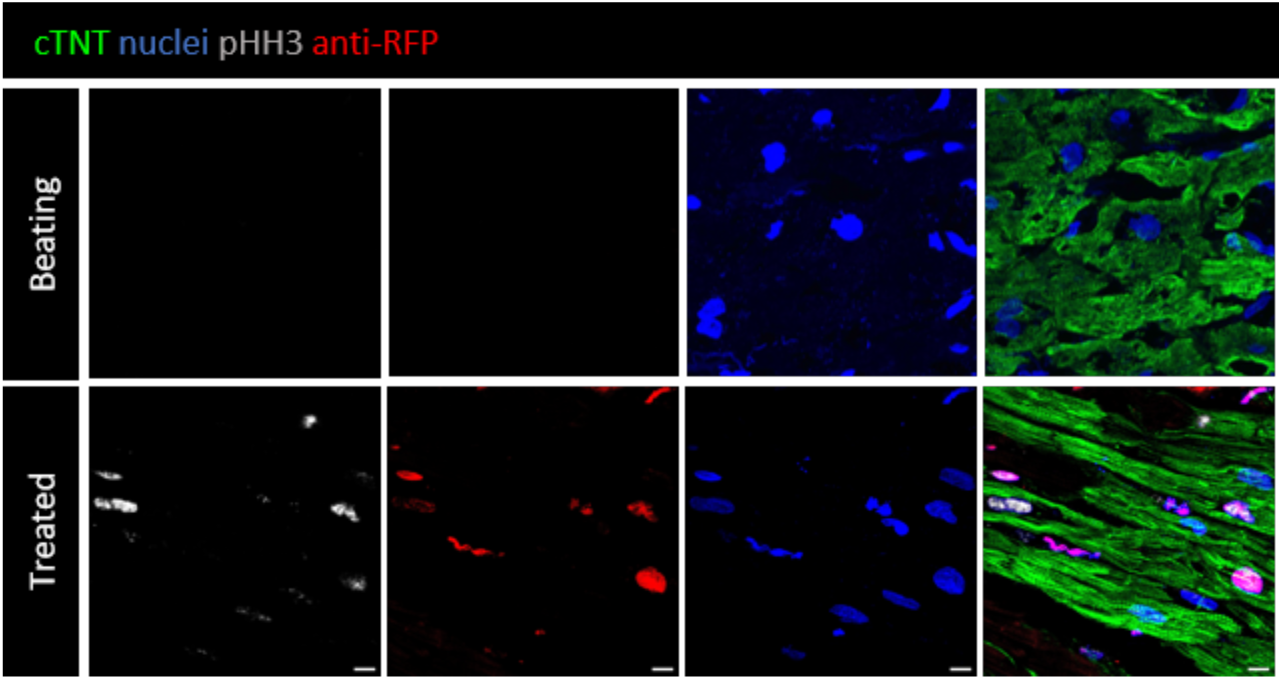
Ex-vivo heart slices transduced with PSAM^4^ shown in lower magnification. Upper row shows a histological staining of an untransduced cardiac slice. Lower row shows a slice transduced with rssAAV6-TnT-nls-TagRFP-2A-PSAM^4^-GlyR treated with varenicline (100 µM) for seven days. Scale bars. 10 µm.

**Fig. S4.**
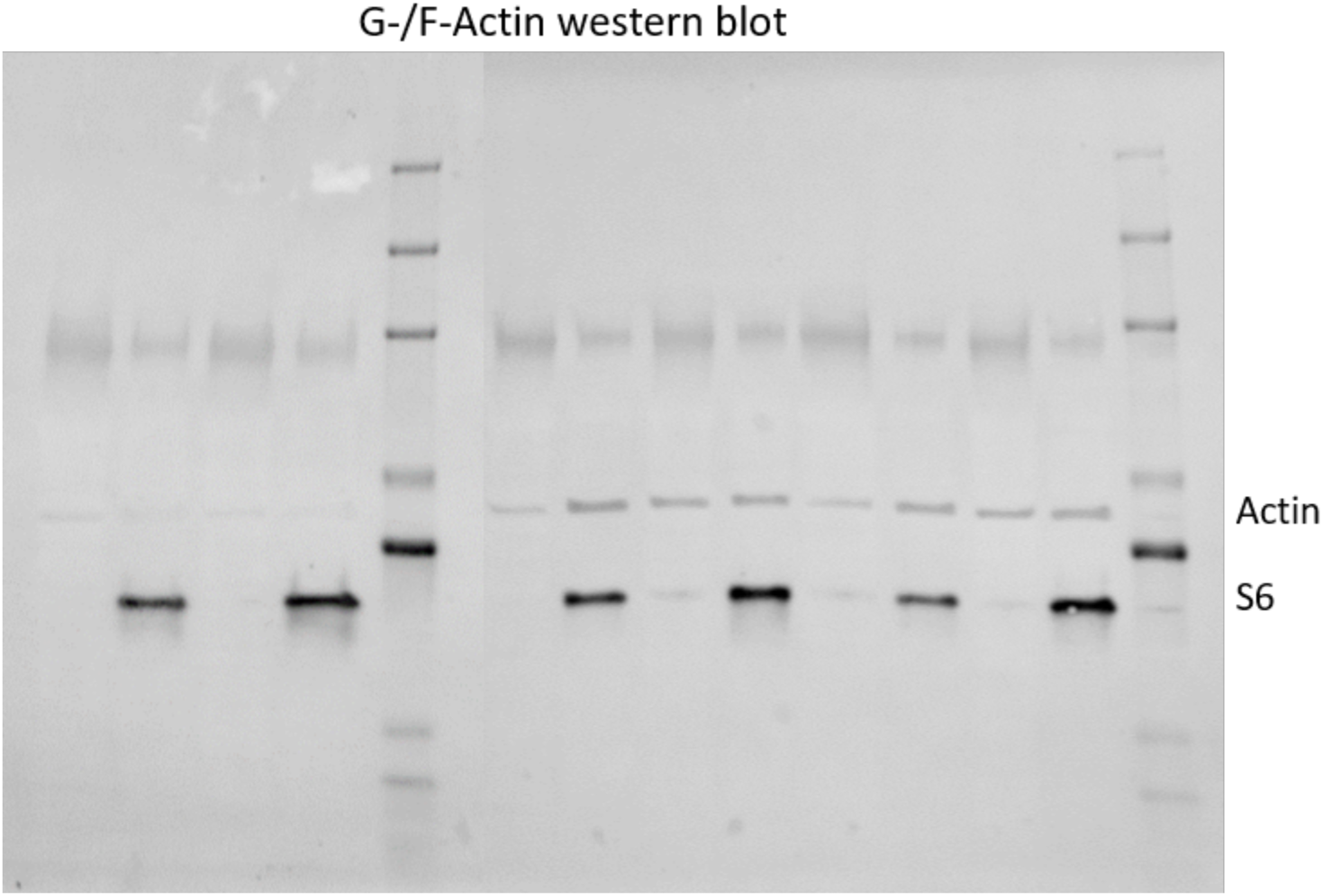
Uncropped Western Blot.

**Fig. S5.**
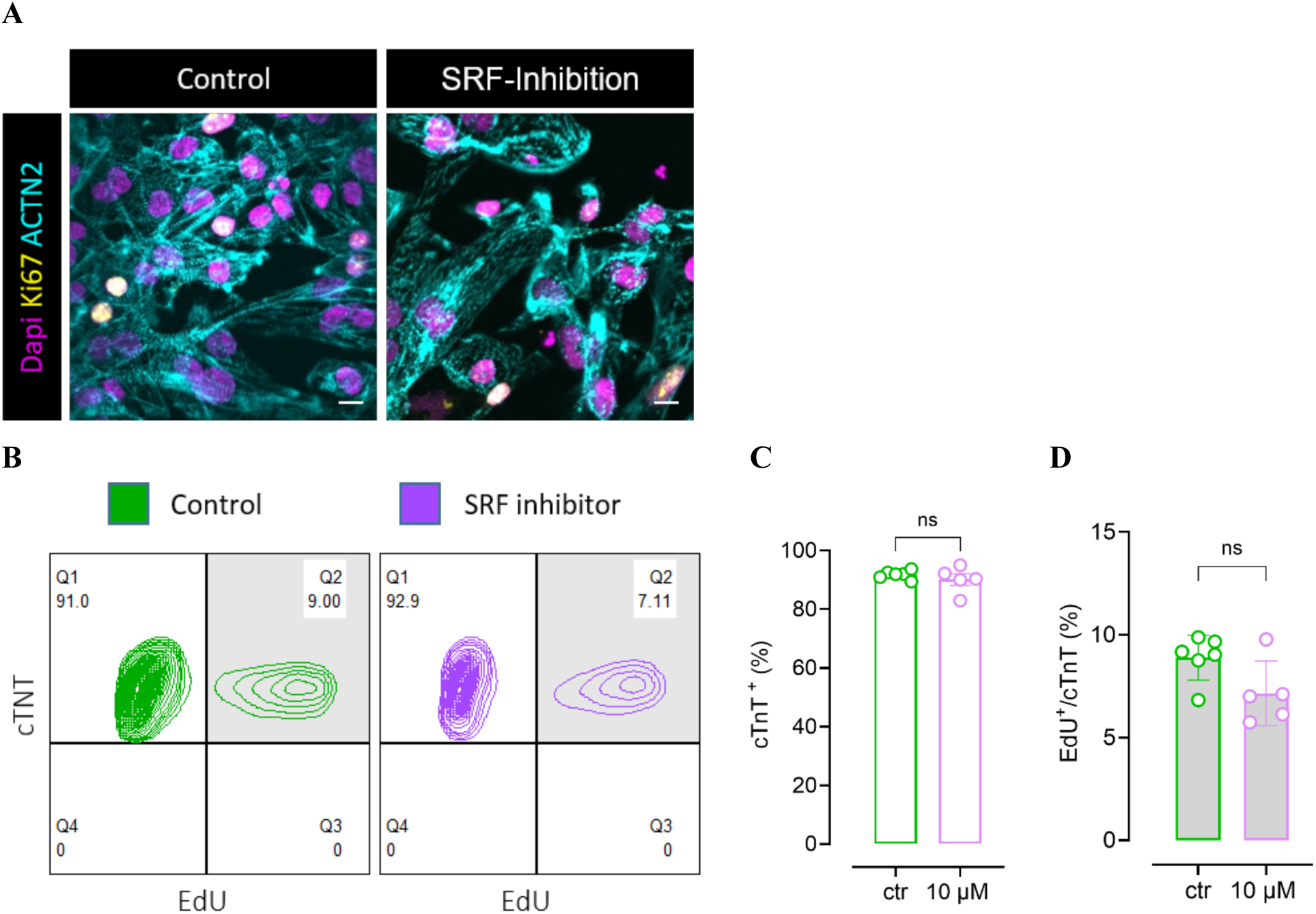
Inhibition of SRF signalling induced sarcomere disassembly but did not promote proliferation. **(A)** Treatment with the SRF inhibitor CCG-203971 (10 µM) led to sarcomere disassembly, as shown by α-actinin staining. (**B)** Representative flow cytometry plots showing EdU incorporation in untreated and CCG-203971-treated cardiomyocytes. (**C and D)** Quantification of (**C**) troponin T (TnT) expression and (**D**) EdU incorporation. Each data point represents one biological replicate from two independent experiments. Scale bars: 10 µm. Statistical analysis was performed using unpaired t-tests.

